# Reproducible patterns of neural activity without attractors in cortical networks

**DOI:** 10.1101/2022.05.24.493230

**Authors:** Domenico Guarino, Anton Filipchuk, Alain Destexhe

## Abstract

Cortical activity often consists of recurring population events of correlated neuronal firing and highly reproducible firing patterns. Because of their resemblance with attractor dynamics, the attractor model prevails today, although it has not been firmly demonstrated. Here, we used a unique dataset, with co-registered two-photon calcium imaging and electron microscopy of the same cortical tissue, to test the central assumption of attractor networks: recurrently active “core” neurons should be strongly interconnected. We report that, contrary to the attractor paradigm, core neurons have fewer weaker connections compared to other neurons. Instead, core neurons funnel the flow of multiple connectivity pathways. Computational models give a mechanistic account of these features showing that distance-dependent connectivity forms converging-diverging motifs and, at their funneling centers, core neurons are found. Thus, reproducible cortical activity and connectivity can be explained without postulating underlying attractor networks but rather by the existence of overlapping information flows.

## Introduction

Before and after birth, every animal has to produce sensorimotor coordination to survive and thrive. This coordination is performed by the whole nervous system, and in particular by cortical population events of correlated neuronal firing^1–6^. Cortical events show reproducible patterns of their participating cells^7,8^ that recur under similar stimulus conditions^7–9^, keep their specificity when reoccurring spontaneously^8–10^, and can be induced by repetitive coactivation^6,11^. Such features seem the outcome of attractor network models, whose dynamics tend to settle into stable, reproducible, activity patterns^3,5,11–13^. Thus, cortical population events are currently used to argue that the overall structure of the mammalian cortex is that of an attractor network^13^.

If reproducible activity patterns are signs of attractor dynamics, then we should be able to find signs of an attractor structure in the cortex. The main structural feature of attractor networks is the existence of strong mutual connections between some units, capable of pulling the network dynamics towards a stored attractor, hence called pattern completion units^12,13^. Although we have so far lacked direct structural evidence, indirect correlation-based functional connectivity^14–16^ has shown strong correlations between neurons having similar evoked responses. However, recent experiments show that repeated optogenetic co-activations do not strengthen synapses, as previously assumed^6,10,13^ rather they increase cell excitability^17^. Also, strong connections are theoretically associated with non-transient attractor states^18^, slowing of network dynamics^19^, and pathological run-away activity^20^.

Here, we tested the currently dominant hypothesis that cortical reproducible activity is the result of attractor networks. We started from the current mapping between theoretical and experimental results. Reproducible population events do not repeat identically. Only a subset of their member cells – called “core” neurons – reliably participates in many event occurrences^7–10,13^. Following the attractor paradigm, core neurons have been assumed to be the biological equivalents of pattern completion units. If it is so, then they should have strong mutual connections. We tested this assumption in the unique MICrONS project^21^ dataset, which offers for the same mouse cortical tissue both a precise mapping of three million synaptic connections – from electron microscopy – and the co-registered spiking activity of more than a hundred neurons – from two-photon calcium imaging. We found that core neurons do not have more abundant, stronger, or mutual connections than other neurons. Instead, we found that core connections are essential for the flow of excitatory activity along the network spanned by reproducible population events.

How is reproducible dynamics possible in a structure showing no evidence of being an attractor network? We provide an alternative mechanistic hypothesis, interlinking a series of known facts. Throughout the cortex, neurons are locally connected in a distance-dependent manner^22–24^, which creates modular networks^25,26^. In modular networks, there is a tendency to form converging-diverging connectivity motifs^27,28^, with central nodes characterised by high-flow connections^29,30^. Forcing the activity flow along same pathways would yield reproducible population events. We tested this chain of hypotheses in balanced spiking network models by varying the spatial range of distance-dependent connectivity. We found that reproducible population events are given only by a limited range of distances, also presenting a large number of converging-diverging modules, having nodes with high-flow connections. We performed the same analysis on the MICrONS dataset obtaining equivalent results.

These findings (i) deepen our understanding of the link between the structure and the dynamics of cortical populations; (ii) open a new view on how to obtain attractor-like behaviours in networks, one that is not based on the strength but on the arrangement of connections; (iii) give mechanistic insight into how the cortex can be ready, at birth, to deliver coordinated sensorimotor facilitation.

## Results

To perform our study, we unfolded the cortical attractor hypothesis as follows: if local portions of the cortex show dynamic signs of storing attractors, like recurring population events, they should also hold structural traces of being attractors. Therefore, we identified core neurons from the population dynamics, and we tested two fundamental attractor-driven assumptions about their connectivity: (i) *synapses between cores are more numerous and/or stronger compared to others*, and (ii) *circuits made by cores involve more recursive connections toward cores*^3,10,12,13^. To perform the tests, we needed information about the *firing* of neurons, to identify events and cores, and the *connections* between those neurons, to perform structural analysis of the circuits in which the cores were embedded. Until recently^28^, this double requirement was impossible to fulfill. The MICrONS project^21^ now gives this opportunity by releasing electron microscopy (EM) and co-registered two-photon calcium imaging datasets of the same cortical tissue (Fig. 1A). Two-photon recordings, and subsequent EM reconstruction, were made in layer 2/3 of the primary visual cortex of an awake mouse presented with static and drifting-oriented noise^33^. The dataset has the duration and temporal resolution (1850 sec, at ~14.8 Hz), and structure (112 cells with both EM reconstruction and two-photon recordings, with 1961 proofread synapses but more than 3 million in total) to perform dynamical and structural analyses. Furthermore, we verified two-photon dataset generality against (N=32) equivalent two-photon recordings from the Allen Brain Observatory^34^.

**Figure 1.**
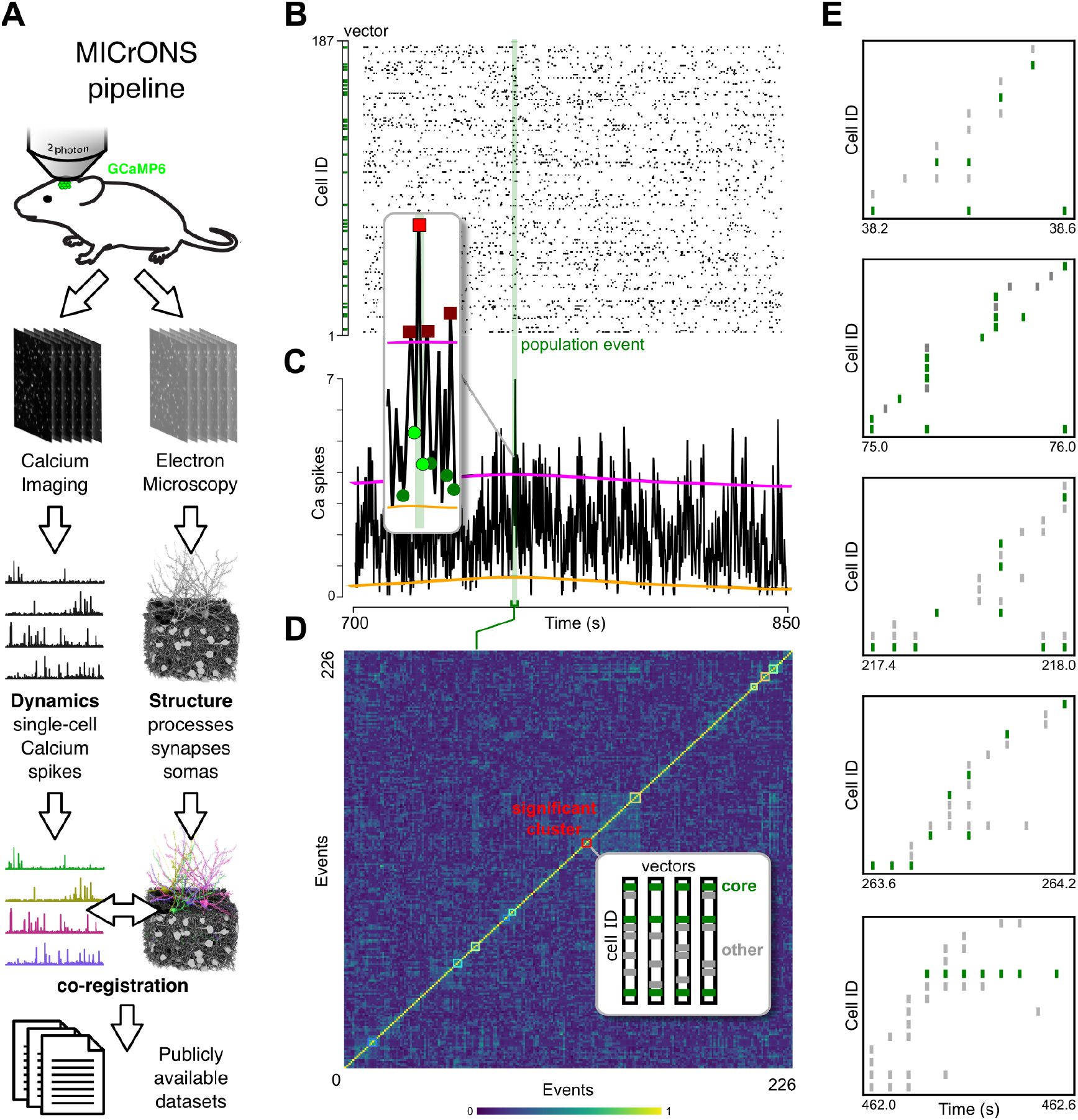
Population events and core neurons. (**A**) Schematic diagram of the MICrONS pipeline. In-vivo calcium imaging and serial EM imaging were performed on the same mouse cortical volume, yielding electrophysiological dynamics and spatial structure of neurons and synapses. An automatized co-registering procedure matched each neuron visible in the calcium imaging movie with a corresponding neuron in the anatomical EM stack. After human proofreading, the data was released to the public. (**B**) Calcium spikes raster plot from awake mice of the MICrONS dataset. Fluctuations in spike frequency indicate synchronous population events (example green shade). (**C**) Population events were quantified as time intervals where the firing rate (black) exceeded a surrogate-based threshold (magenta) above firing baseline (orange). *Inset*. The subthreshold minima (green points) before and after each peak (red squares) were the beginning and end of each event. (**D**) Each event was assigned a population binary vector (B, participating cells in green). Each population binary vector was correlated with all others (color bar, Pearson’s correlation) and clustered based on their similarity. Population events sharing neurons beyond a surrogate-based threshold were identified (colored squares, non-significant clusters were omitted), their autocorrelation computed, and the cells participating in their events beyond a threshold were labeled as “cores” (*inset*). (**E**) Example events from the cluster highlighted in D. Cells are displayed sorted by time of occurrence.

To identify reproducible spiking patterns, and their core neurons, we expanded a previously described method^6–12,31^ for the analysis of two-photon calcium imaging rasters (Fig. 1B). Briefly, population events were identified as peaks of synchronous firing deviating significantly from a surrogate-based threshold (Fig. 1C). The neurons firing within an event were listed in an event vector (Fig. 1B, *left*). Pearson’s correlation between event vectors was then used to gather events into clusters (Fig. 1D). Clusters with significant pattern reproducibility – the correlation between event vectors within the cluster – were then identified as those beyond a surrogate-based threshold. Finally, core neurons were singled out as those participating in the events of significant clusters (Fig. 1D, inset) beyond a threshold justified below. In addition, we confirmed core identities using functional correlation^25^ (Fig. S1). The MICrONS two-photon recording presented 226 significant events, and 10 clusters of significant reproducibility (>0.25), with 35 core neurons, reliably occurring across events (see example events in Fig. 1E). Events number, duration, and pattern reproducibility were similarly distributed in the MICrONS and Allen Brain Observatory datasets (Fig. S2).

### The connectivity of core neurons is not consistent with pattern completion units

Once cores were singled out, we proceeded with testing the attractor-driven assumptions. For the first – *synapses between cores are more numerous, and/or stronger, compared to others* – we took the number and volume of postsynaptic spines as a proxy for synaptic efficacy^35–37^. We started by assessing whether core neurons made (or received) on average more numerous or bigger spines compared to any other neuron (irrespective of whether them being or not part of the two-photon imaged dataset to have good statistical power, see Methods). Core neurons did not make more numerous (Fig. 2A) nor bigger spines (Fig. 2B) with any neuron, compared to other non-core neurons. Then we turned to the (core-to-core, core-to-other, other-to-core, other-to-other) connectivity of neurons participating in significantly reproducible events, hence limited to both EM and two-photon imaged neurons. Given the total of imaged cells and their proofread spines, the count of spines per core was underpowered. Nonetheless, we used the current proofread dataset to study core and non-core synapse distributions, adopting two inclusive solutions: (i) we lowered the threshold for core identification to a minimum (participation to 60% of the events), and (ii) we did not differentiate spines based on their point of contact (e.g. axo-somatic, axo-dendritic, axo-axonic, etc). Core neurons presented no connections among themselves, and a higher probability of having connections to and from non-cores compared to others (Fig. 2C). In addition, the postsynaptic spine volumes made by core neurons were not bigger than those made by other neurons (Fig. 2D).

**Figure 2.**
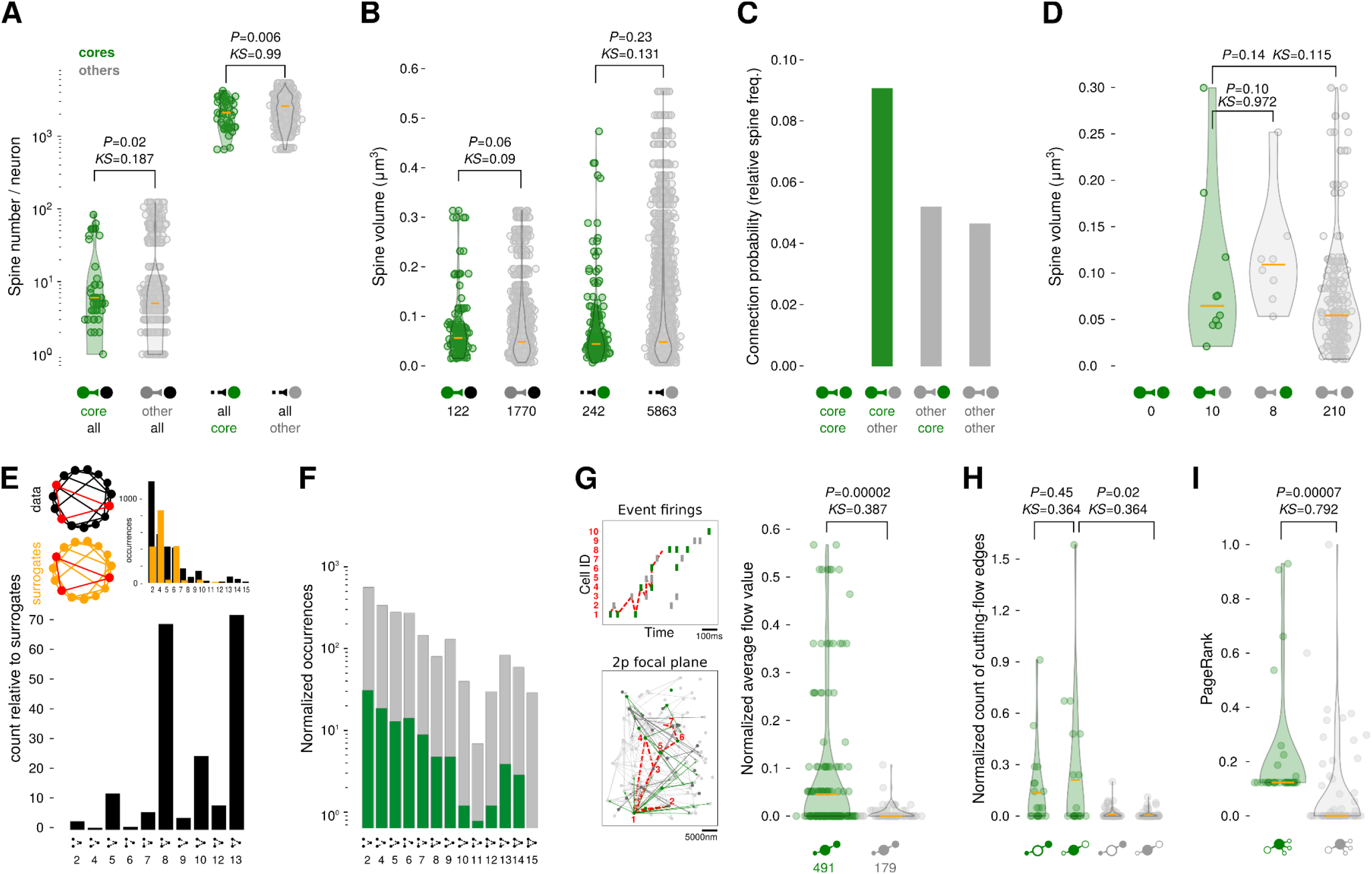
Core neurons support the flow of population events. (**A**) Number of postsynaptic spines per neuron made by 2-photon imaged cores (green) and other non-core (gray) neurons to all (black) neurons within the EM volume (*left*), and received from neurons within and beyond the volume (*right*). Cores made significantly more connections locally (but with poor KS sample size) and received significantly fewer connections from within and beyond the volume. (**B**) Postsynaptic (proofread) spine volume made by all cores and other neurons to all neurons within the EM volume (*left*), and received from neurons within the volume (*right*). There was no significant difference between spine volumes. (**C**) Connection probability for core and other non-core neurons. Note the absence of core-to-core synapses. (**D**) Postsynaptic spine volume distributions of cores and others were not significantly different. (**E**) Connectivity motifs distribution for all two-photon recorded neurons (*top:* occurrences count). (**F**) Normalized motif occurrence distribution for cores and other neurons. Both made a mixture of mutual and non-mutual connections. (**G**) Event flow analysis. (*Left*) The sequence of active cells in each population event (*top*) identifies a subnetwork (*below*). By following all possible pathways between active cells in sequence, we retrieve each cell’s flow capacity. (*Right*) Cores had significantly more (normalized) flow capacity. (**H**) Removing connections having cores as targets (green, *right*) stopped significantly more event flows than connections targeting non-cores (gray, *right*). Cores were equally sources or targets of cutting-flow edges (green *left* vs *right*, non-significant difference). (**I**) Core neurons had significantly higher PageRank scores than non-cores.

For the second assumption to test – *circuits made by cores involve more recursive connections towards cores* – we performed several graph theory analyses, considering cells as *nodes* and synapses as *edges* of the network^26^. To get a global picture of mutual connectivity, we looked at the assortativity *r* – the proportion of mutual edges in a directed graph. We found *r*=-0.08, characteristic of non-mutually connected networks. To get a detailed picture, we looked at the distribution of motifs – statistically significant connectivity patterns considering groups of three cells^38^. To understand the significance of motif occurrences, we used the ratio of real vs 100 surrogate degree-matched networks^39^. The ratio followed a known distribution^39^ showing an abundance of three-cell mutual motifs (Fig. 2E). However, looking at the motif distribution as a function of cores and other neurons across all population events of each cluster, we found that mutual, as well as convergent and divergent, motifs were not specific to cores (Fig. 2F).

Even if not having direct mutual connections, core mutual activity could have been sustained by indirect synaptic feedback, via secondary or looped pathways, abundant enough to ensure that there were cliques of pattern completion units in traditional attractor network terms^12^. These possibilities entailed the existence of more edges, paths, and cycles involving cores than non-cores. We measured the number of cliques, the number and length of shortest paths, and cycles between cores or non-cores. None gave a significant advantage to cores (Fig. S3).

### Core neurons support the flow of population events

The relatively large number of edges towards and from cores suggested that core neurons could be central to many cortical pathways. However, simple measures of centrality (degree, betweenness, and hub score^26^) gave no significant difference between cores and other neurons (Fig. S4). But these measures are only concerned with the static – instantaneous – existence of edges between dynamically characterized nodes. Population event dynamics are instead not instantaneous, but rather a sequence of cell activations in time (Fig. 2G, left *top*, dashed red line) and space (*bottom*, dashed red line). The *flow* algorithm^40^, by weighting all possible edges between pairs of active cells in sequence, retrieves each cell’s capacity to sustain that specific sequence. Cores had significantly more flow capacity than others (Fig. 2G, *right*). The counterpart of flow is *cut* – the minimal set of edges that, if removed, would disconnect the sequence. Cores were on average significantly more targets of such edges (Fig. 2H). This result was confirmed by measuring the *PageRank^26^* – which assigns scores to a neuron proportional to the scores of the neurons it is connected *by*, thus being a weighted measure of convergence within each event. Core neurons had significantly higher PageRank scores than others (Fig. 2I).

Altogether, there were no more numerous or stronger or more recursive connections between cores. Instead, we found that cores had high sequence flow capacity, were the target of cutting edges, and had high PageRank. Sequence flow capacity tells the specificity of the circuits targeting cores. The PageRank score tells how relevant the neurons converging on cores are. Together, the two measures indicate that the convergent connections funnel incoming activities towards cores, suggesting that this could be the source for the reproducibility of the events they belong to.

### The locality of cortical connections yields reproducible cortical activity patterns

What is the connectivity principle giving reproducible population events, and creating circuits with cores as structural funnels, in networks showing no evidence of being attractor networks? Based on the strong connectivity assumption of attractor networks, models have been proposed where strong recursive paths were imposed on randomly connected networks by either additional circuits or increased synaptic strength between arbitrarily selected neurons^13,41,42^. These models obtain synchronous firing events corresponding to the activation of recursive sub-population(s). However, the explanatory and predictive power of these models is diminished by their ad-hoc solutions^2^, and we have presented evidence casting doubts on this assumption.

Alternatively, it is known that distance-dependent connectivity rules, without imposing strong recursive connections, produce weakly recursive networks that spontaneously and irregularly transition between periods of higher and lower firing rates^43–49^. Building on these results, we present a mechanistic hypothesis: distance-dependent connectivity rules make hierarchically modular networks, where each module has multiple converging paths funneling towards multiplexing “core” units. We wove this hypothesis by interlinking known facts. Distance-dependent connectivity creates modules by allowing mostly short-range and few long-range connections^43,45,50^. Such modular structure has been shown to occur between the cells of mouse sensory cortices^25^ and the human cortex^47^. Crucially, in hierarchically modular networks, each module tends to form a convergent-divergent connectivity motif, known as “bow-tie”^27,28^. Connections in such a motif are arranged to funnel long and short-range contributions toward the central nodes, characterized by high max-flow min-cut values^29,30^. In a spiking network, these neurons would become more frequently active, without recurrent pathways.

We tested our hypothesis in models by systematically varying the range of connection distances between neurons and verifying that (i) distance-dependent connectivity creates hierarchically modular networks, with (ii) bow-tie modules, (iii) having high-flow nodes, (iv) which overlap with the dynamically identified cores of reproducible population events. To perform the tests, we developed a biologically-informed, yet minimal, model of a cortical network of 10000 excitatory regular-spiking, and 2500 inhibitory fast-spiking, conductance-based Adaptive Exponential Integrate-and-Fire model neurons^51^. In building our model, we established connections for each neuron by randomly sampling its neighborhood with probability exponentially decreasing for increasing distances (Fig. 3A). We chose parameters such that connectivity loosely followed the EM distribution of connection distances (Fig. S5). Within these limits, we systematically varied the connectivity range to study the necessary and sufficient conditions to have hierarchical modularity, bow-tie structures, flow, and population events as those observed in-vivo. For model neuron dynamics, we used already published parameters from the mouse cortex^44,45^. The model was driven by external independent Poisson spike generators (proportional in number to Fig. 2A synapse counts).

**Figure 3.**
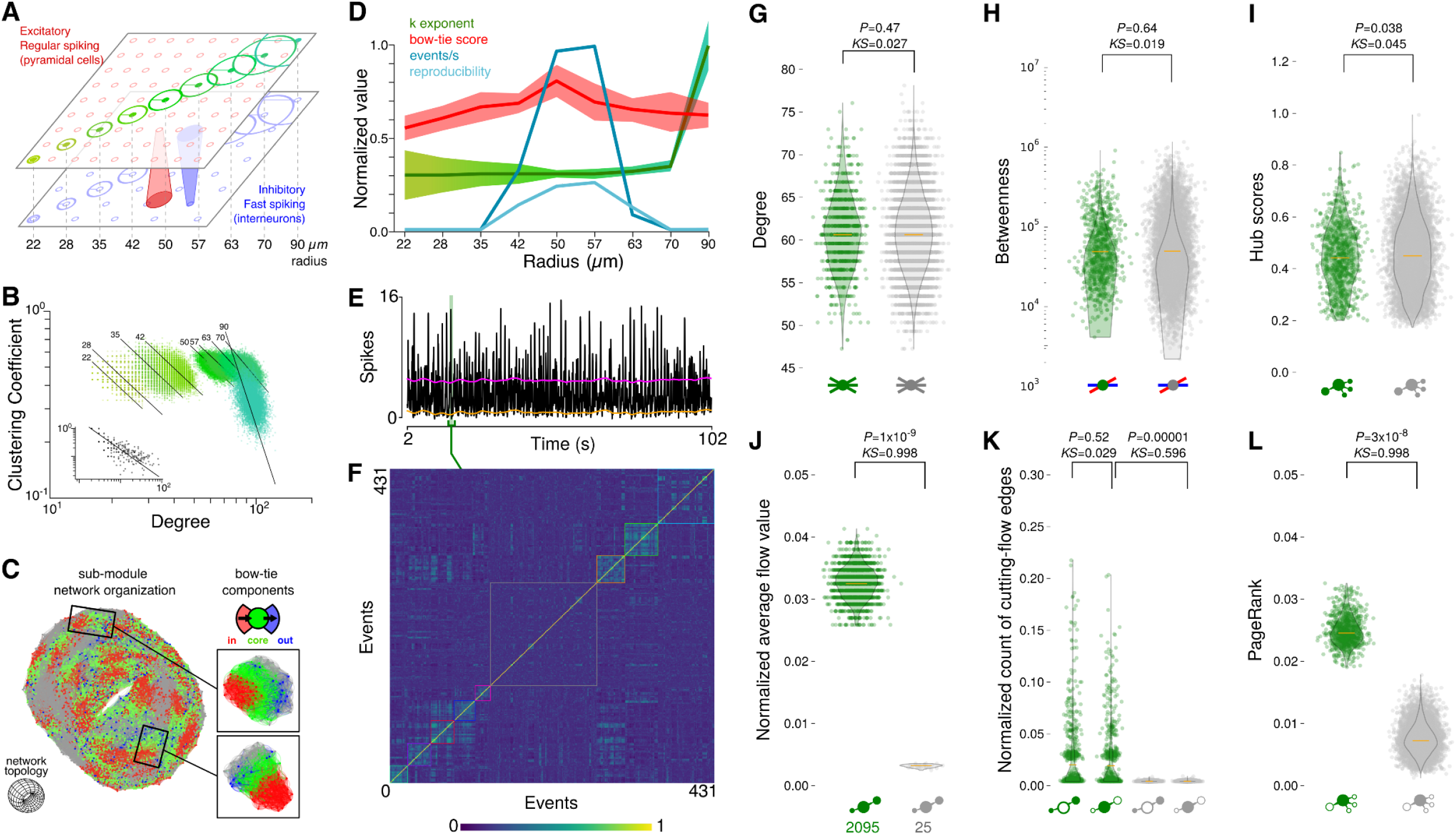
Distance-dependent connectivity creates high-flow core units. (**A**) The model was made of excitatory (red) and inhibitory (blue) neurons on a 2D grid of ~1000μm (10μm step). Within biological limits, 9 connection range radiuses were tested. (**B**) All connection ranges resulted in hierarchically modular networks – i.e. having power-law relationship between degree and local clustering coefficient of their neurons – (inset, MICrONS data). (**C**) For each range, some network modules could be further analyzed as bow-tie motifs (top right) with in- (red), core (green), and out- (blue) components. Two example modules (right). An example network (50μm radius) is shown (left) using the same color code (gray, units not assigned to any module). Its toroidal topology (idealization, bottom left) is customary by design, to avoid bordering conditions. (**D**) Summary scores for the different models. The power-law fitting (green shades as in **A**) shows constant modularity for all models (except for large ranges). Bow-tie score (red) peaked for mid-ranges (42-57 μm). The normalized number of events (dark blue) and reproducibility scores (light blue) also peaked in the same range. (**E**-**L**) Example core analysis for an example network (50μm radius). (**E**) Population events were quantified as in Fig. 1A. (**F**) Population binary vectors were correlated with all others and clustered as in Fig. 1C (gray squares identify non-significant clusters). (**G**) The distribution of cores’ number of connections was not different from other neurons. (**H**) Cores were not on more pathways than others. (**I**) Cores were not preferentially connected to highly connected neurons. (**J**) Core neurons sustained significantly more than others the flow of the event sequences they participated in. (**K**) Removing core connections from the network graph stopped more event sequence flows than others. (**L**) Core neurons had significantly higher PageRank scores than non-cores.

### Distance-dependent connectivity creates hierarchically modular networks

Within the biologically realistic limits just described, we checked that several connection distances are sufficient and necessary to produce modular networks (Fig. 3B). A network is hierarchically modular^25,26^ if, for each of its neurons, there is a power-law relationship between the local clustering coefficient *C(d)* – the ratio of connected neighbor pairs over the total number of neighbor pairs – and their degree *d*, i.e. such that fitting an exponential function to the local clustering and degree relationship yields a proportion *C(d) ~ d^-k^*, with exponent *k-l*^25^. Overall, this means that the network is made of many low-degree locally well-connected nodes and few high-degree less-locally connected nodes, resulting in a scale-free modular architecture. We found that all connectivity ranges produced hierarchically modular networks, with exponent *k* close to 1 (*k*=[1.02, 1.3]), as found experimentally^25^ (and Fig. 3B, black inset for the MICrONS data *k*=1.05). Distance-dependent connectivity is also necessary for hierarchical modularity. Beyond the physiological limits (>200μm), modularity disappeared from the model, due to the increased reach of connections and the consequent lack of local clustering, and a version of the same network, with randomized connectivity preserving node degree, presented no modularity.

### Distance-dependent connectivity creates bow-tie modules

For the ranges that yield hierarchically modular networks close to the MICrONS data, we inspected the organization of connectivity within each module. Given that our data and models are directed, and that our focus is on the flow of information, we identified modules as the most frequented subgraphs spanned by (hundreds of) random walks over directed edges^52^. Once modules were identified, the convergent-divergent bow-tie organization^27,28^ of each module was assessed by finding its largest subgraph component (Fig. 3C, inset green), then verifying the existence of an in-component – projecting to the largest (Fig. 3C, red) – and an out-component – receiving from the largest (Fig 3C, blue). If all three components were present, the module was counted as bow-tie, otherwise as simple module (Fig 3C, gray). To be able to compare bow-tie organization across connectivity ranges, we defined a bow-tie modularity score as the ratio of bow-tie labeled over the total number of modules (bow-tie + simple). The bow-tie score was above 0.5 for all connectivity ranges but it rose for ranges around 50 μm (topping at 50 μm, with 0.82±0.3, Fig 3D red). In comparison, the MICrONS graph had a lower 0.14±0.1 bow-tie score, but randomly reassigning increasing portions of each node’s targets to either closer or farther nodes (keeping node degree), to mimic a procedure similar to our models, resulted in a degradation of the bow-tie score comparable to that observed in the models when departing from the 50 μm range (Fig. S6A-C).

### Distance-dependent connectivity creates high-flow core units

For all ranges that yielded hierarchically modular networks, we tested the presence of population events and cores with flow statistics comparable to experimental results. Over the 9 ranges, only 4 produced synchronous irregular population events with reproducible patterns and cores with funneling flows. These corresponded to the networks with hierarchical modularity exponent *k*~1, and the highest bow-tie modularity (Fig. 3D), supporting the theoretical hypothesis. The results of an example model with those characteristics is given in Fig. 3E-L. The model produced synchronous irregular dynamics, whose content switched between population events characterized by different event vectors. On these events, we applied the same core analysis as above (Fig. 3F), resulting in 431 significant events. Of these events, 273 made 8 clusters of significant reproducibility (>0.2), with 896 core neurons.

The distance-dependent connection strategy was sufficient to get a similar arrangement of circuits found in the MICrONS dataset. Model cores had degree, betweenness, and hub scores distributions not significantly different from other neurons (Fig. 3G-I). As in that dataset, the model presented no evidence of more mutual connectivity via indirect synaptic feedback for cores (Fig. S7). But, importantly, compared to others, model cores had significantly higher sequence flow capacity (Fig. 3J), they were significantly more the sources and targets of cutting edges (Fig. 3K), and they had significantly higher PageRank (Fig. 3L), matching the structural-dynamical measurements of real core neurons.

The distance-dependent connection strategy was also necessary. We took the above network and randomly rewired all connections, preserving each neuron’s node degree. This randomly rewired network gave rise to population events, but no reproducible pattern was identified, and consequently no cores (Fig. S8). The reason is that by increasing the connectivity range, the number of connections and the number of paths increases such that the funneling disappears^53^. A completely random network has an infinite connectivity range and no reproducibility. Trying to rescue event reproducibility in random networks, by increasing input correlations, leads to population-wide oscillations far from biologically realistic regimes (Fig. S8D).

### Dynamical cores overlap with structural cores

Finally, we checked the consistency of our hypothesis chain by looking at the overlap between dynamically-identified core neurons – those reliably participating in multiple clustered population events – and structurally-identified core neurons – those repetitively found in bow-tie modules (Fig. 4A). In the models, dynamical core neurons matched the central neurons of bow-tie modules with an average of 9.7% (min: 1.7%, max: 36.7%, see Fig. 4B right). The MICrONS data had a higher match, with an average of 21.2% (min: 0.1%, max: 66.2%, see Fig. 4A left). Two considerations concern these values. First, given the number of neurons (model: 10000, data: 112) and cores (e.g. model: 896, data: 35), the number of possible combinations is so high that the percentages are significant (Fig. 4B, red line is the max intersection between all possible combinations of sizes equal to the core sets, see Methods). Second, different strategies were used to find structural and dynamical cores. Structural cores are high-convergence nodes resulting only from random walks along the locally available edges. Instead, dynamical core neurons receive orders of magnitude more connections from the global than the local network (Fig. 2A). Therefore, they are co-active across events clustered for reasons depending also on non-local sources (external drive in the model, cortico-cortical and/or thalamic common inputs in the experimental data).

**Figure 4.**
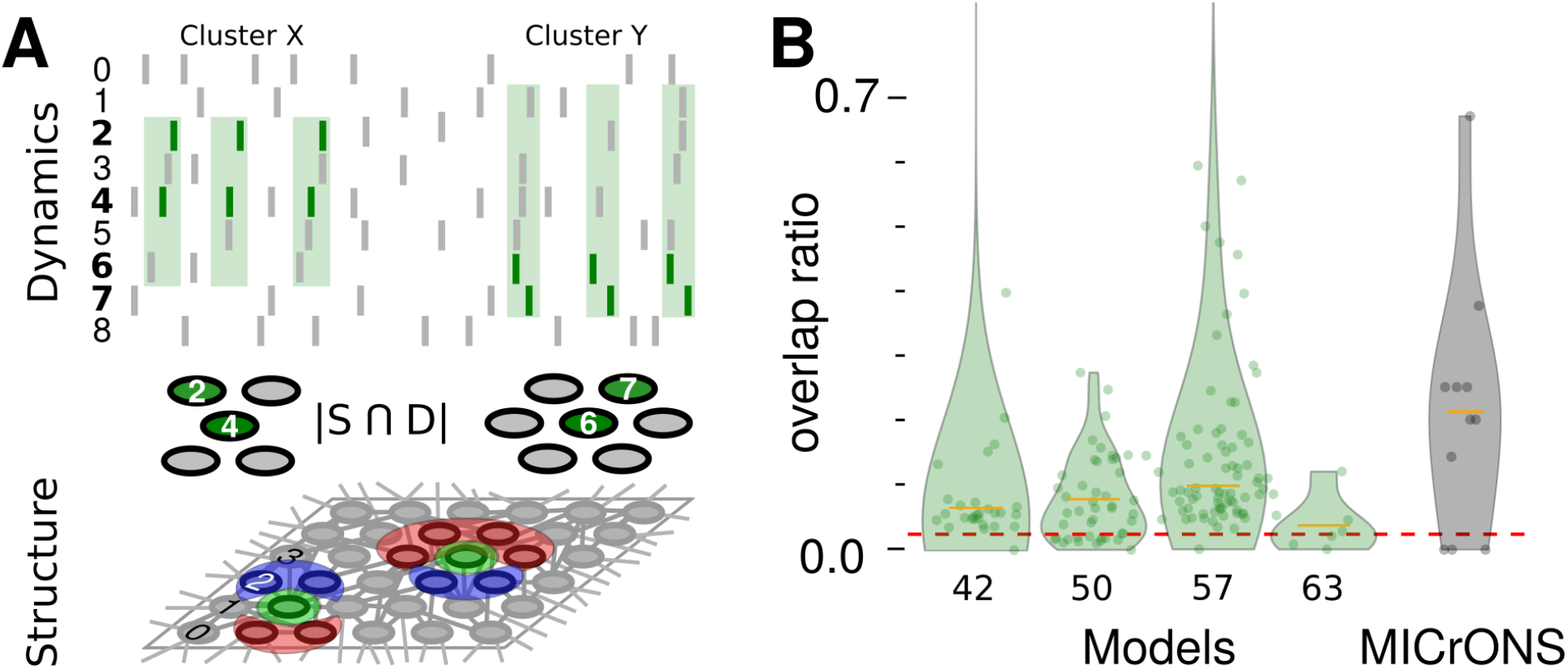
Dynamical cores overlap with structural cores. (**A**) Dynamical (top) and structural (bottom) clusters of cells were identified with different strategies. Dynamical clusters (top, groups of green shaded bands) were found by recurring co-activation, structural clusters (bottom, groups of red-green-blue bow-ties) by recurring random paths. Once the two types of clusters were found, we checked their consistency by measuring the overlap between them, counting how many dynamically-identified recurrently active cells were also structurally-identified as centers of bow-tie modules. (**B**) The overlap ratio – size of the intersection of dynamical and structural core sets – of models (left, green data points and violin plots for each model yielding cores [42, 50, 57, 63 μm radius], three outliers not shown) and MICrONS data (right, grey data points and violin plot) is on average higher than the significance threshold (red dashed line, analytic significance obtained by calculating all possible values of the overlap ratio under rearrangements of the data, *p*=0.024, see Methods).

## Discussion

Uniformity and modularity have been used to argue in favor of an attractor-like structure of the cortex^2,3,13,54^. Accordingly, the role of cortical networks in cognition would be understood as transformations between attractor representations, where the identity of, and connections between, specific neurons can be safely regarded as details^54^. However, here we looked at local portions of the cortex showing dynamic signs of being attractors – like reproducible population events and cores – and searched for structural traces of attractors – like enhanced connectivity between core neurons – we found none. When we explored the space of distance-dependent connectivity in models, we found that reproducible population events are the results of a converging-diverging, not attractor-like, organization of connections.

There are two major implications of our results. The first is a missing element in the mechanics introduced by Hebb. All animals need the sensorimotor coordination of cell assemblies, which in Hebb’s description are forged by learning^1,3^. Then, how could the cortex deliver coordinated activity at birth without any prior learning? From day one, before any experience-driven learning, the cortical system must be wired to have groups capable of recurrent co-activation with (some) functional specificity. Without motor facilitation and consequent action, all post-action evaluation mechanisms would not operate on cortical synapses^55^ to shape cell assemblies. The models we built within biological plausibility, supported by the experimental data analysis, suggest how distance-dependent connectivity can jump-start the cortex into being able to deliver coordinated facilitation from day one. Onto a cortex pre-wired for distance-dependence, Hebbian learning will fine carve functional specificity, potentiating and depressing connections. We have both theoretical and experimental indications that distance-dependent connectivity is genetically encoded^56^, and that sensorimotor functional maps are created already during embryonic development^57^. Interestingly, Hebb^1^ hinted to such a possibility in a note (page 121).

The second implication strengthen the link between microscopic and macroscopic scales for the spontaneous and evoked activity across global brain states. Spontaneous activity was found to define the realm of possible evoked responses in cortex^58^ in the anesthetized state. But in the awake state, spontaneous activity was later found to be orthogonal to the evoked, when also behavior is involved^59^. Here we can add a mechanistic element in line with the literature about this apparent paradox^8^. In our results, the clusters of cells found using only their structural connectivity only partially overlap with the clusters found using cell activities (Fig. 4). Dynamical cores are reliably co-active across events clustered for reasons that can only partially be reduced to their local connectivity. In fact, we found that core neurons receive orders of magnitude more connections from the global network than what they return (Fig. 2A left vs right). Thus, during anesthesia, when global excitability is reduced, the local structural connectivity may become more dominant^60,61^, explaining the uniformity of spontaneous and evoked responses^8,58^.

There are several limitations to our study that need to be taken into account when evaluating the present and future efforts: (i) a single mouse dataset necessarily gives a reduced and biased picture, (ii) its two-photon/EM sample size is still limited, and (iii) sensory cortical areas are less likely to have an attractor-like structure compared to association areas^12,62^. To overcome these limitations, it can be mentioned that the coordinated EM/two-photon approach will generate more datasets, improving the present picture. In addition, we supply a model built using very conservative assumptions, which closely mimics structural and dynamical experimental features.

Future directions of work should deepen our understanding of the connectivity structure and better examine the nature of the incoming activity in the network. The generic distance-dependent strategy we adopted is susceptible to varying sampling distributions beyond exponential to explore more complex functions^46,49^, which can be parametrized to represent different cortical areas. In our models, the external drive (Poisson generators) represents the input from the rest of the global network. It needs to be variate, as increasing input correlations introduce global oscillations. However, apart from distance-dependent connectivity, we took no other parameter into account. What would happen if additional features were added to the organization of connectivity, or to the input correlations, is currently under study. For example, Filipchuk et al.^8^ recorded in the auditory cortex, where local connections are not only hierarchically modular but also tonotopically organized, and incoming activity has correlations inherent to the stimulus.

If cell assemblies have to sustain thought processes and behaviors, they need to be flexible but also reliable. These requirements made the attractor model appealing for its resilience to perturbations. But when closely inspected, the cortical structure does not seem to hold any sign of being an attractor. A valid alternative can be a distance-dependent arrangement of connections yielding, prior to any learning, a versatile substrate to shape cell assemblies.

## Acknowledgments

This work has been funded by EC Human Brain Project (HBP, grant agreement no. H2020-945539). We express our gratitude to the MICrONS project and the Allen Brain Institute for the efforts they make and their commitment to publicly releasing their datasets, without which this study would have not been possible. We would like to thank Adrienne Fairhall, Yves Frègnac, Manuel Mameli, and Federico Tesler, for stimulating discussions and critically reading the manuscript. The mouse drawing in Fig. 1 is offered by Ethan Tyler and Lex Kravitz.

## Author contributions

Conceptualization: D.G., A.F., A.D.

Methodology: D.G., A.F.

Software: D.G.

Validation: D.G., A.F.

Investigation: D.G., A.F., A.D.

Visualization: D.G., A.F.

Funding acquisition: A.D.

Project administration: A.D.

Supervision: A.D.

Writing – original draft: D.G., A.F., A.D.

Writing – review & editing: D.G., A.F., A.D.

## Materials and Methods

We performed the same structural and dynamical analyses over data collected in mouse cortex and in biologically constrained conductance-based spiking models.

### MICrONS data source

The MICrONS project (https://www.microns-explorer.org/) made available, through its website, two datasets, corresponding to the first two phases of the project. The first phase, the one we used, has electron microscopy and functional imaging of the same cortical patch of V1, in a 400 × 400 × 200 μm volume with the superficial surface of the volume at the border of L1 and L2/3, approximately 100μm below the pia. Here follows a brief description of the procedures, and we refer the interested reader to Dorkenwald et al. 2022. **Two-Photon imaging**. Functional imaging was performed in a transgenic mouse expressing fluorescent GCaMP6f from the Jackson Laboratories (jax.org) and the Allen Brain Institute (see next section on the Allen data source). The complete description of mouse transgenic line, surgical procedures, implants, two-photon microscope features, stimulation and recording setups, and protocols were already published (Dorkenwald et al. 2022). One important feature and the main difference with the Allen Brain Observatory two-photon dataset is the recording acquisition frame rate, here at ~14.8 Hz, resulting in time-frame windows of ~67 ms (the Allen dataset has double this resolution, ~30 Hz). **Stimuli**. The mouse was head-restrained but could walk on a treadmill during imaging. Visual stimuli were 30 one-minute trials of a colored-noise interspersed with periods of coherent motion of oriented noise. Each one-minute trial contained 16 stationary-moving-stationary blocks, with a different direction presented in each block, pseudorandomly-ordered. **Electron microscopy**. After functional imaging, the mouse was perfused with fixant agents, dissected, and 200 μm coronal slices of the V1 area, interested by two-photon sessions, were cut. Serial sections of 40 nm thickness were cut from the slices and prepared for transmission electron microscopy. Custom modifications to the setup and camera allowed fields-of-view as large as (13 μm^2^) at 3.58 nm resolution. Data were acquired in blocks, aligned through multiple steps, and defects manually detected. Somas, synaptic cleft, and spine detection and partnering were manually performed on a (387 clefts) subset for training and testing machine learning libraries. The rest were performed by machine learning libraries. Finally, cells identified in the two-photon imaging max projection were manually co-registered with EM-identified somas.

### Allen data source

The Allen Institute made available, through its Brain Observatory, a large dataset of neuron responses from several visual cortical areas and layers, using mouse transgenic lines expressing a Cre/Tet-dependent fluorescent calcium indicator (GCaMP6f), which allows the recording of calcium influx associated with neural activity. The complete description of mouse transgenic lines, surgical procedures, behavior training, implants, two-photon microscope features, stimulation and recording setups, and protocols were already published (see Allen VisualCoding_Overview whitepaper in the references). Here we briefly provide the relevant information in the context of our work. **Cell type and experiment selection**. The Allen Brain Observatory contains data collected in several cell types of mouse visual cortices (see Allen VisualCoding_Genotyping whitepaper). Cell-type specific expression of GCaMP6f was conferred by the Cre recombinase and tetracycline-controlled transactivator protein (tTA) under the control of the Camk2a promoter with the Ai93 and Ai94 reporters. To maximize the number of recorded neurons across layers, we selected all Cre drivers specific for pyramidal neurons and layers. Considering that each layer of mouse V1 is roughly 100um thick (Hage et al. 2022), we were able to design cell-type and layer-specific queries which resulted in the selection of the following experiment ids (as of 2019/09/10). We selected layer 2/3 planes, in order to have a dataset comparable to the one from the MICrONS project (query: http://observatory.brain-map.org/visualcoding/search/overview?area=VISp&imaging_depth=175,185,195,200,205,225,250,265,275,276,285,300&tld1name=Slc17a7-IRES2-Cre,Cux2-CreERT2.Emx1-IRES-Cre), selected experiment ids: [502115959, 501574836, 502205092, 502608215, 510514474, 503109347, 510517131, 524691284, 645413759, 660513003, 704298735, 702934964, 712178511, 526504941, 528402271, 540684467, 545446482, 561312435, 596584192, 647155122, 652094901, 652842572, 653122667, 653125130, 657080632, 661328410, 661437140, 663485329, 679702884, 680156911, 683257169, 688678766] (with exp: 501271265 removed due to https://github.com/AllenInstitute/AllenSDK/issues/66). A total of 32 experiments were selected and downloaded as NWB files. Each experiment contained pre-processed calcium indicator fluorescence movies (512×512 pixels, i.e. 400 μm cortical field of view, sampled at 30Hz), corrected for pixel leakage during scanning, ROI filtered, and neuropil subtracted. Each resulting NWB file consisted of an experiment session (~90 min) at a given location (cortical area and depth).

All original code of this study has been deposited at https://github.com/dguarino/Guarino-Filipchuk-Destexhe and is publicly available as of the date of publication. DOIs are listed in the key resources table. The MICrONS project dataset is available at: https://www.microns-explorer.org/phase1. The Allen Brain Observatory dataset is available at: http://observatory.brain-map.org. The calls to retrieve the Allen data are listed in the Materials and Methods section. An interactive Jupyter notebook is accessible on Binder: https://mybinder.org/v2/gh/dguarino/Guarino-Filipchuk-Destexhe/HEAD. The code to run all simulations (complete with Dockerfile to recreate the environment) is available at https://github.com/dguarino/Guarino-Filipchuk-Destexhe-simulations. Any additional information required to reanalyze the data reported in this paper is available from the lead contact upon request.

### Calcium spike timing extraction

To infer the action potential timings from the calcium indicator fluorescence of the Allen Brain data source, we used the python library OASIS (Friedrich et al. 2017). This library for spike inference, which is based on an active-set method to infer spikes, also has routines for automatic estimation of its parameters. The library is available at https://github.com/j-friedrich/OASIS. As parameters for the deconvolve function, we used *penalty*=1, and we used the automatic estimation of fluorescence variation from the data (parameter *g*). The resulting spike timings obtained with these parameters were verified for adherence to the example deconvoluted data provided by the Allen Brain Observatory ipython notebooks.

### Dynamical analysis

For each data source, we applied the same dynamical analysis. The analysis of synchronous irregular population events has already been developed in MATLAB™ and described in Filipchuk et al. 2022 (itself derived from Miller et al. 2014, and going back to Prut et al. 1998), please refer to it for an in-depth description. We reimplemented all analysis steps in Python and tested them against the original analysis. Briefly, single cell activity was described as a cell binary vector, where the number of elements was equal to the *T*, number of time frames during the recording (frame duration was ~30ms). A value of 1 was assigned at the time frames corresponding to the onset of each spike, and the cell vector was zero otherwise. The number of cell vectors was equal to the total number of recorded neurons in the field of view, *N_e_*, resulting in a *TxN_e_* spiketrain matrix where columns corresponded to the time frames and rows to the neurons. Each column was summed to give the number of neurons coactive at every time frame from which the instantaneous population firing rate could be deduced. As is visible from sample raster plots (Fig. 1A), there were periods of time where spikes were much more synchronized across the population, which could be interpreted as population events, departing from the fluctuations of an asynchronous population spiking process. To identify whether the population events were above chance coincidence, and determine their exact start and end, we applied the following algorithm. (**1**) We created 100 surrogate *TxN_e_* spiketrain matrices by reshuffling each neuron’s inter-spike interval. The average 99 percentile across time frames was then calculated and a firing rate threshold for event significance was obtained by adding this average value to a local baseline estimate that aimed at correcting slow fluctuations of background firing rate. The baseline estimate was generated using an asymmetric least squares smoothing algorithm from Eilers and Boelens (2005), with parameters *p=0.01* for asymmetry and *λ*=*10^8^* for smoothness. (**2**) Experimental population firing rate trace was smoothed using Savitzky-Golay filtering, with order=3 and frame window=7, to get rid of non-essential peaks. Local maxima for the smoothed curve that were above the previously defined threshold were retained (Fig. 1B, inset, red square). The adjacent local minima around each of the local maxima were identified (Fig. 1B, inset, green circles). Their time frames were taken as the beginnings and ends of population events (Fig. 1A, limits of the green vertical band). (**3**) The time interval between the beginning and end of each population event defined its duration. We described each population event as a binary vector of length equal to the total number of neurons in the recorded field of view (*N_e_*). We initialized all vector’s elements to *0*, and then we indicated the participation of a cell to the event, spiking at least once over its duration, with a coordinated *1* (Fig. 1A, left). (**4**) Pearson’s correlation matrix for ongoing and evoked events was constructed by computing the Pearson correlation coefficient between all binary vectors. In plotting the matrix, the color reflected the degree of correlation with a colormap ranging from 0 to 1. (**5**) Population events sharing a substantial number of neurons, i.e. having a significant degree of correlation, were clustered together (using agglomerative linkage clustering with the farthest distance based on correlation metric, from the python *scipy.clustering* package), and the correlation matrix could be permuted according to the clustering order (Fig. 1C). (**6**) The correlation between population events might be simply due to the finite number of recorded cells. We, therefore, defined a significance threshold for intra-cluster correlation, established using surrogate events. We generated *100*x*Ne* surrogate event vectors (with the same number of cells per event of the originals, but randomly chosen cells, without replacement). We then performed correlation-based clustering (as in point 5) of the resulting surrogate clusters, and we used the 95 percentile of correlation coefficient of these clusters as a correlation threshold. (**7**) To quantify the pattern reproducibility of events within a cluster, i.e. the number of cells shared across events in a cluster, the correlation was calculated across all events of each cluster. (**8**) Core cells were identified as those participating in a range of thresholds (from 55%, barely above no difference in participation, to 95%) of the events within each cluster. Relaxing this requirement to allow for a more participative definition of cores changed the statistics but not the trends presented in Fig. 2. Once identified, we verified that the average firing rate of each core was higher within than that outside its events, discarding those with unspecific firing. In developing our dynamical analysis for the MICrONS data, we used the notebook example on the Allen Institute (https://github.com/AllenInstitute/MicronsBinder/blob/master/notebooks/vignette_analysis/function/structure_function_analysis.ipynb). The whole analysis workflow code, is available on GitHub (https://github.com/dguarino/Guarino-Filipchuk-Destexhe), including also an interactive Jupyter Notebook version on Binder for faster reproducibility (https://mybinder.org/v2/gh/dguarino/Guarino-Filipchuk-Destexhe/HEAD).

### Structural analysis

We used the data tables published by the MICrONS project on their website and the referenced publications (see above). **Postsynaptic spine number and volume analysis**. We selected the subset of proofread synapses having as soma root id those identified and co-registered in the two-photon dataset. After the dynamical analysis was performed (see above), core neuron ids for each event were available and used to query the synapse table, and postsynaptic spine volumes were collected by identified type (core or non-core). The four possible combinations (core-core, core-other, other-core, other-other) were plotted in Fig. 2. Given that different cores participated in different clusters, the number of cores and others differed, and each core (or non-core) neuron could potentially be connected to all other cores (or non-cores) we reported in Fig. 2A the normalized number of spines. Differences in significance and effect size were established using Kruskal-Wallis and Kolmogorov-Smirnov distance, since the distributions were non-normal, as reported below in the statistical methods. **Synaptic functional efficacy analysis**. To estimate functional connectivity, we adopted the same method described in Sadovsky and MacLean (2013). Briefly, an adjacency matrix representing a graph for each experiment was created with nodes being all recorded cells, and directed edges formed according to a single frame lagged correlation (frame duration: 67.4ms) and weighted according to the number of spikes (see supplementary figure 1). **Graph theory analyses**. An adjacency matrix was built by coordinating pre- and post-soma root ids. All graph analyses reported in Fig. 2 were performed using the python version of the library *iGraph* (Csárdi and Nepusz, 2006). The iGraph library allows motif identification with either 3 or 4 nodes. We used 3-motifs to be able to compare our results with other publications and with the online MICrONS ipython notebooks, and also to make computations of the 100 surrogate graphs run in a reasonable time. The comparison of core-based cycles and other-based cycles was performed over an adjacency list and using a function that recursively checked over the list of neighbors starting with each core (or non-core) cell id. In developing our structural analysis for the MICrONS data, we used the notebook example on the Allen Institute GitHub page (https://github.com/AllenInstitute/MicronsBinder/blob/master/notebooks/intro/MostSynapsesInAndOut.ipynb). The whole analysis workflow code, including also an interactive Jupyter Notebook version on Binder, is available (see above). **InfoMap analysis**. In order to identify structural modules in the MICrONS data and the models, which are directed network, we used a random walk approach described by Rosvall and Bergstrom (2008). We also used the same algorithm to identify, within each module, the largest connected component (supposedly the center of a bow-tie connectivity motif) and the in-/out-components to the central one.

### Neuron model

All simulated networks were composed of n=12500 (excitatory/inhibitory neuron ratio of 4:1, i.e. 10000 excitatory and 2500 inhibitory) conductance-based adaptive exponential integrate-and-fire (AdExIF) point neurons. The set of equations describing the membrane potential (1) and spike adaptation current (2) of a neuron are

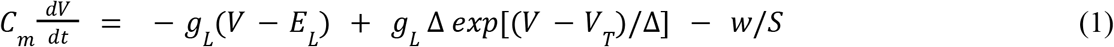

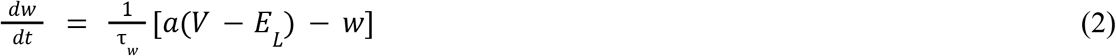

with *C_m_* being the membrane capacitance (nF), *g_L_* the resting (leak) conductance (μS), *E_L_* the resting (leak) potential (mV, which is also equal to the reset value after spike), Δ the steepness of the exponential approach to the threshold (mV), *V_T_* the spike threshold, and *S* membrane area (μm^2^). When *V* reaches the threshold, a spike is emitted and *V* is instantaneously reset and clamped to the reset value during a refractory period (ms). *w* is the adaptation variable, with time constant *τ_w_* (ms), and the dynamics of adaptation is given by parameter *a* (in μS). At each spike, *w* is incremented by a value *b* (in nA), which regulates the strength of adaptation. An experimentally informed parameter set was chosen (from Zerlaut et al. 2011) and kept fixed, see Table 1, here below.

**Table 1.**
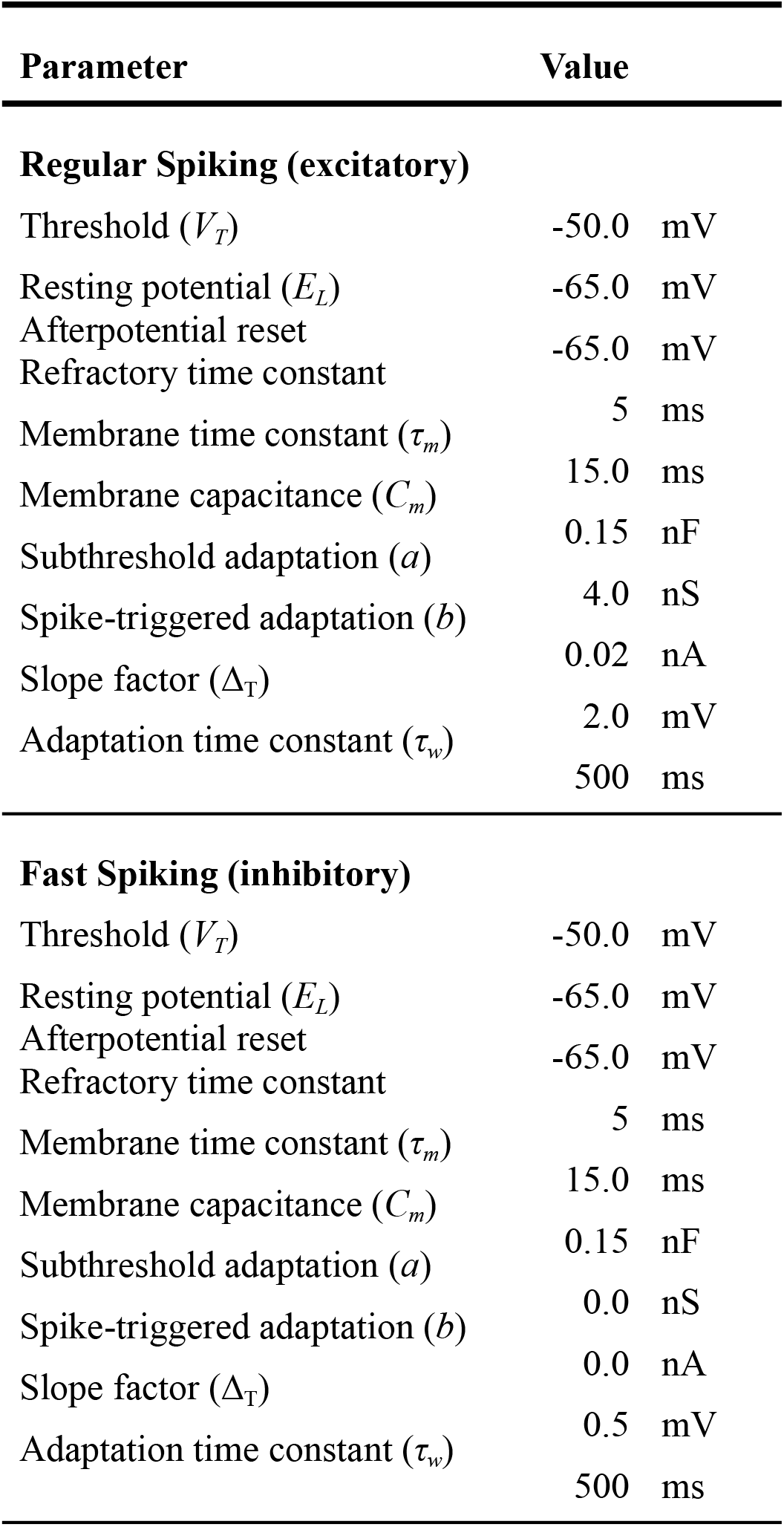
Parameters of the AdEx Integrate-and-Fire model.

| Parameter | Value |
| --- | --- |
| **Regular Spiking (excitatory)** |  |
| Threshold ($V_T$) | -50.0 mV |
| Resting potential ($E_L$) | -65.0 mV |
| Afterpotential reset | -65.0 mV |
| Refractory time constant | 5 ms |
| Membrane time constant ($\tau_m$) | 15.0 ms |
| Membrane capacitance ($C_m$) | 0.15 nF |
| Subthreshold adaptation ($a$) | 4.0 nS |
| Spike-triggered adaptation ($b$) | 0.02 nA |
| Slope factor ($\Delta_T$) | 2.0 mV |
| Adaptation time constant ($\tau_w$) | 500 ms |
| **Fast Spiking (inhibitory)** |  |
| Threshold ($V_T$) | -50.0 mV |
| Resting potential ($E_L$) | -65.0 mV |
| Afterpotential reset | -65.0 mV |
| Refractory time constant | 5 ms |
| Membrane time constant ($\tau_m$) | 15.0 ms |
| Membrane capacitance ($C_m$) | 0.15 nF |
| Subthreshold adaptation ($a$) | 0.0 nS |
| Spike-triggered adaptation ($b$) | 0.0 nA |
| Slope factor ($\Delta_T$) | 0.5 mV |
| Adaptation time constant ($\tau_w$) | 500 ms |

### Synapse model

The synaptic connections between neurons were modeled as transient conductance changes. The synaptic time course was modeled as an alpha function with a fast rise followed by an exponential decay:

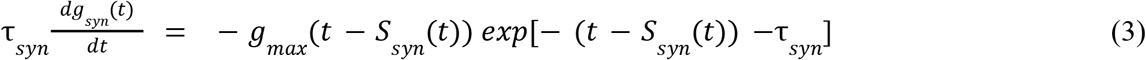

where *syn∈{exc, inh}, S_syn_(t)*=*δ(t – t_k_)* are the *k* incoming synaptic spike trains where *i∈{1, N}* refers to presynaptic neurons and k to the different spike times of these neurons. The synaptic time constants were chosen to be *τ_exc_=3* ms and *τ_inh_=7* ms for excitation and inhibition respectively. The reversal potentials were *E_exc_*=0 mV and *E_inh_*=-80 mV. The synaptic weights *g_syn_(t)* were set to *g_exc_*=1 nS for the excitatory conductance, and a balance of *g_inh_*=5 nS unless stated otherwise. We used a constant delay of 0.5 ms, equal for all connections.

### Network architecture

We simulated a microscope focal plane area of ~1000 μm^2^ as a 2D-layer-like network (with periodic boundary conditions to avoid size effects). Neurons were arranged on a grid meant to represent ~10 μm separation between neurons. Every neuron was sparsely connected with the rest of the network with a connection probability drawn from distance-dependent probability distribution. Each neuron *i* was connected with another neuron *j* with a probability *p_ij_*, depending on their distance *r_ij_*, according to an exponential profile:

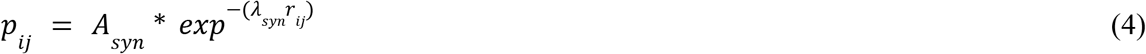

where *A_syn_* is the amplitude of the exponential, varying the number of neurons sampled in the chosen radius *r_ij_*. The amplitudes *A_syn_* were set to have the number of connected neurons roughly match the distributions described by Katzel et al. 2011, Markram et al. 2015, and Seeman et al. 2018, considering the topological distribution and the excitatory/inhibitory neuron ratio (*A_exc-exc_*=14, *A_exc-inh_*=24, *A_inh-exc_*=14, *A_inh-inh_*=24), resulting in a degree of ~75 connections per neuron. The decay controller ***λ**_syn_* was set to ***λ**_exc_*=1.2 and ***λ**_inh_*=1.5. The seed to the random number generator responsible for the sampling of the connections was systematically varied, producing N=20 different networks, but following the same type of exponentially decaying distributions. In the tests for the relationships between distance-dependence and flow measures, we systematically varied the amplitudes, lambdas (and correspondingly synaptic weights) to keep the conductance balance constant across network configurations. The arrangement of connections was tested with respect to changes in synaptic weights (spplementary figure S9).

| radius<br>($\mu\text{m}$) | power-law<br>fit ($k$) | $R^2$ | normalized<br>bow-tie / tot<br>communities | units /<br>community | population<br>events / s | pattern<br>reproducibility |
| --- | --- | --- | --- | --- | --- | --- |
| 22 | 1.02 | -3.45 | 0.56±0.2 | 42.6±0.6 | – | – |
| 28 | 1.02 | -0.65 | 0.61±0.2 | 49.2±0.5 | – | – |
| 35 | 1.03 | 0.15 | 0.68±0.3 | 68.3±0.3 | – | – |
| 42 | 1.03 | 0.10 | 0.70±0.1 | 82.6±0.4 | 1.00 | 0.14 |
| 50 | 1.03 | 0.01 | 0.82±0.3 | 113.1±0.2 | 2.92 | 0.25 |
| 57 | 1.03 | 0.06 | 0.71±0.4 | 128.0±0.3 | 2.99 | 0.27 |
| 63 | 1.20 | 0.04 | 0.65±0.2 | 153.3±0.1 | 0.24 | 0.12 |
| 70 | 1.30 | 0.03 | 0.62±0.5 | 177.0±0.4 | – | – |
| 90 | 3.34 | -1.57 | 0.60±0.2 | 253.7±0.5 | – | – |
| data | 1.05 | -10.5 | 0.17±0.1 | 22.5±0.6 | 1.21 | 0.18 |

### Drive

An external population of Poisson drivers was connected to both the RS and FS populations, with uniform random probability drawn from the half-open interval [0,0.01), with a fixed synaptic weight of 0.5 nS.

### Simulator

All simulations were performed using the NEST simulator (Diesmann and Gewaltig 2001) controlled through the PyNN interface (Davison et al. 2009).

### Statistical Analysis – cores and spines

The data released during phase 1 by the MICrONS project concerned one mouse. This limit was our major concern while designing our study. However, our unit of statistical analysis was the number of events. Events are produced by the same cortical tissue overall but by different cells and connections. This implies that we did not set out to measure the same outcome multiple times on the same animal – which would entail an assay variability. Instead, we set out to measure different (quasi-)independent outcomes – which is inherent biological variability (the high level of interconnectivity does not make it completely independent). This is why, in the currently released MICrONS dataset, we consider as statistically legitimate our sample size of n=226 events, i.e. the number of independent observations of our unit of analysis, under a single experimental condition. The goal of our study was hypothesis testing, therefore we designed each analysis as a statistical hypothesis test to produce an exact statistical significance level (*P-value*) with an appropriate effect size. Concerning the attractor features for cores vs non-cores, they are taken to exist if the following hold in the specific direction (one-tailed direction): (1) Core neurons have more numerous synapses between themselves, or towards others, compared to those of non-cores. (2) Spines with pre-synaptic cores and post-synaptic cores or non-cores are bigger than spines with pre-synaptic non-cores. **FIG. 2AB**. We started by assessing whether core neurons made (or received) on average more numerous or bigger spines compared to any other neuron (irrespective of them being or not part of the two-photon imaged dataset). The required sample size, common to both points above (but here calculated for the spine volume), can be computed by considering the following. The study group (“cores”) is compared to the rest of the population (“all”). The primary endpoint is a continuous value (synaptic spine volume, μm^3^). The anticipated mean of the “others” is 0.05±0.2 (based on the recorded average). Estimating a higher average for cores of 0.1, with a type I error probability (false positive rate) of α=0.05, a type II error probability (false negative rate) of β=0.25 (corresponding to a 75% power), the resulting sample size for core spines is 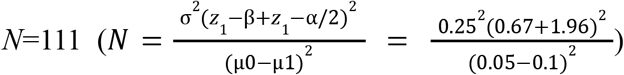. And the smallest count of spines (core-to-all, Fig. 2B, left) is *N*=122. **FIG. 2CD**. We then turned to the (core-to-core, core-to-other, other-to-core, other-to-other) connectivity of neurons participating in significantly reproducible events, hence limited to both EM and two-photon imaged neurons. The required sample size, common to both points above, can be computed by considering the following. The study group (“cores”) is compared to the rest of the population (non-cores, or “others”). The primary endpoint is a continuous value (synaptic spine volume, μm^3^). The anticipated mean of the “others” is 0.02±0.12 (based on the recorded average). Estimating a higher average for cores of 0.08, with a type I error probability (false positive rate) of α=0.05, even with a type II error probability (false negative rate) of β=0.3 (corresponding to a modest power of 70%), the resulting sample size for cores is *N* = 254. The actual number of two-photon recorded cells is *N*=112, but the number of cores per event varies between 3 and 8. Therefore, in the current status of the MICrONS project, the spine count for cores is underpowered. In order to provide an estimate, we considered four aspects: (a) given that the spine count for each cluster of events with its cores and others is taken independently, many spines will be considered in a repeated measure fashion; (b) the maximum value of the independent variable (spine volume) is >10 times its minimum value (respectively 0.35 and 0.02 μm^3^), allowing a large space of possible values; (c) we lowered the threshold for core identification to a minimum (participation to 60% of the events); (d) we did not differentiate spines based on their point of contact (e.g. axo-somatic, axo-dendritic, axo-axonic, etc).

### Statistical Analysis – dynamical vs structural cores

After the bow-tie analysis on modules (above) identified structural cores, we needed to know their overlap with the dynamical cores (the bow-tie procedure involving random jumps across the graph, its results may slightly vary with different calls). To compare dynamical and structural cores, we used the intersection between each set of dynamical (D) and set of structural vector of cores by the total number of dynamical cores (having always the largest number of elements): (D ∩ S) / D. Given the large number of elements in both the MICrONS data and (even more) the model, we computed an estimation of the significance level by permutation test. Model: 9571 cores in 47 clusters (328±0.2) out of 10000 neurons (*p*=~10^13656^). MICrONS: 35 cores in 11 clusters (3.6±0.32) out of 112 neurons (*p*=5.24×10^14^, see Fig. S6D). We computed analytically the significance threshold by calculating all possible values of the overlap ratio under rearrangements of the data. The *p*-value (common to both structural and dynamical cores for simplicity) was the probability of intersection of size equal to the average size observed between all combinations of size as the average size of structural core sets, *X_s_*, and all combinations of size as the average size of dynamic core sets, *X_d_*, where the overlap is 21.2% (as in the MICrONS data). If the observed occurrences were lower than those achieved by random combinations with replacement we considered the occurrences significant. Thus, the *p*-value results as the probability of two combinations sampled from the same population to have an intersection of size *m*, |*X_s_* ∩ *X_d_*| = *m* (with *m*=2 on average for the MICrONS data) is

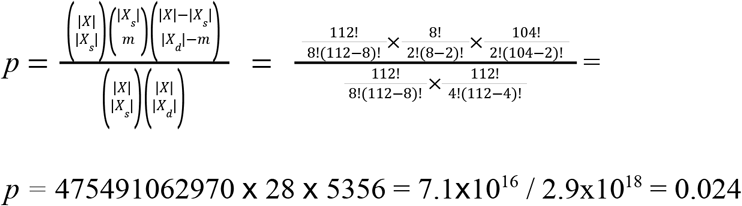

which is what is reported in the figure 4, panel D, red dashed line.

### Statistical Analysis – general

For all statistical tests, to evaluate the differences between distributions resulting from our measurements of the nominal variables (spine volumes, 1-lag correlations, degrees, betweenness, …) we considered that: the nature of the outcome was continuous, the number of groups was larger than 2 (unless specified, or with unequal measurements), the groups were independent (cores and non-cores groups do not share cells). We tested the distributions’ normality using the D’Agostino and Pearson’s test. For normally distributed variables, we used the One-way ANOVA test. For non-normally distributed variables, we adopted the Kruskal-Wallis test. For some variables, the sample size was large (beyond 100 samples), therefore even small variations could be labeled as significant by the above tests. In these cases, a quantitative measure of the effect size was required. As a measure of effect size, we adopted the Kolmogorov-Smirnov test difference, which is more robust than Cohen’s delta for non-normal distributions (Wilcox and Keselman 2003). For all these tests, we used the Python *scipy* library.

## Supplemental Informations

**Fig. S1.**
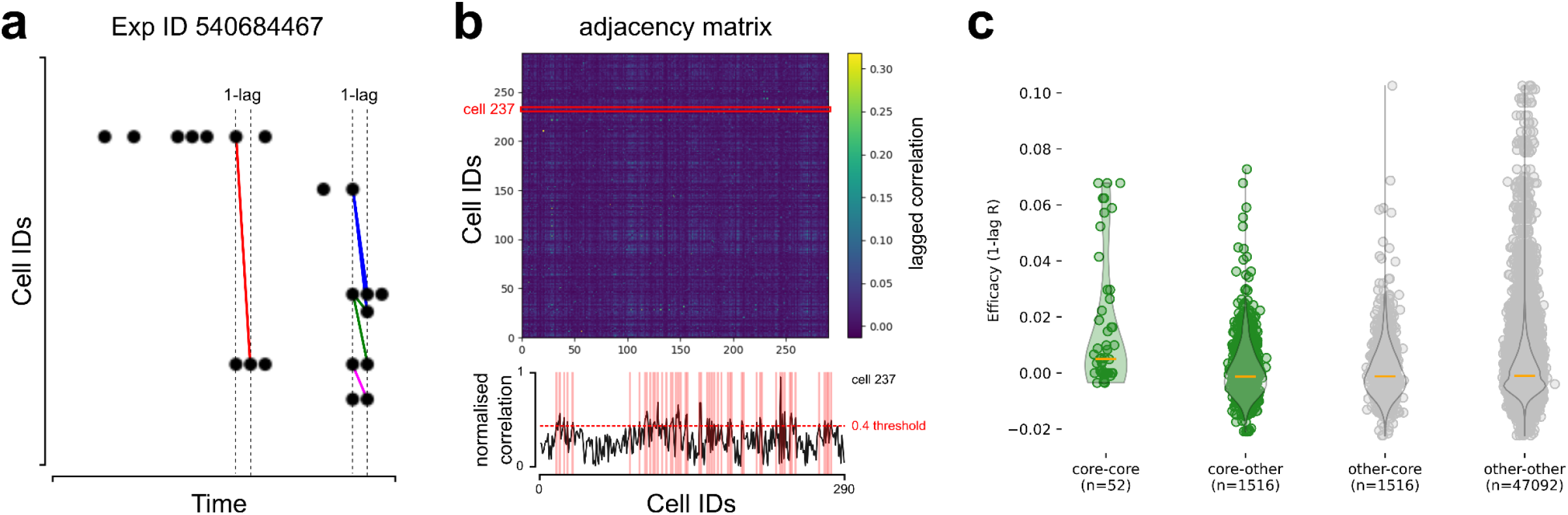
Core functional correlation. (**A**) Detail from the Allen Brain Observatory experiment ID 540684467 raster plot showing calcium spikes (black circles). Each row contains the calcium spikes for a cell. The minimal interval between two spikes is the two-photon imaging time resolution (30 Hz for the Allen Brain dataset). This interval represents the 1-lag correlation interval to identify cells potentially connected. In the figure, only some cells with 1-lag correlated firing were joined by colored segments to improve legibility. (**B**) The vector of 1-lag correlations for each cell is collected into a matrix (top). Weak correlations (threshold=0.4) were not considered for further analysis (bottom, correlations for cell id 237). (**C**) Core neurons are those sharing (non significantly) more high functional correlation across events compared to others.

**Fig. S2.**
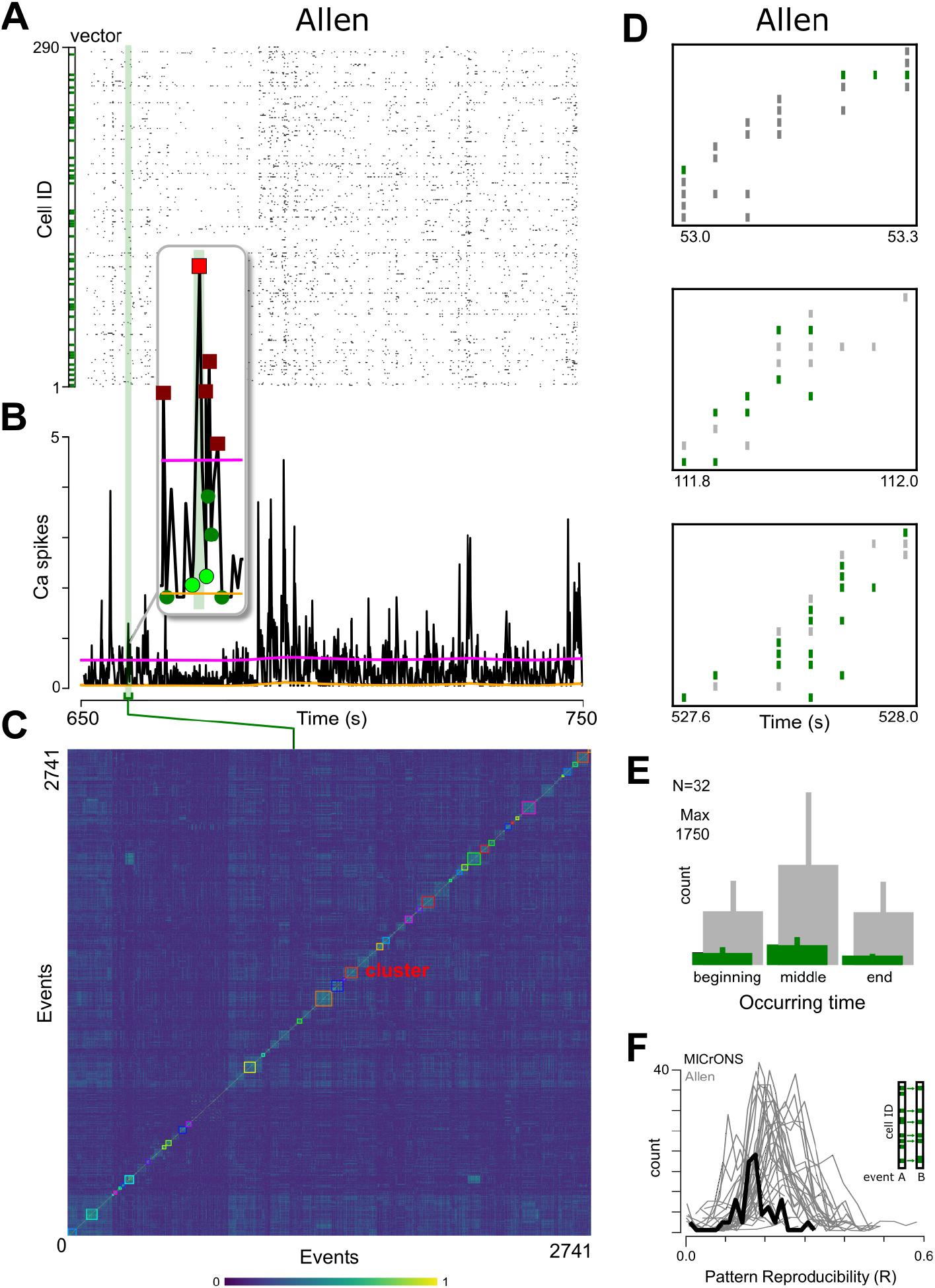
Comparison of MICrONS and Allen Brain Observatory events. (**A**-**E**) As for Fig. 1, Fig. 1. (**A**) Calcium spikes raster plot from an example experiment (ID: 540684467). (**B**) Population events. (**C**) Clustered matrix of correlations between population vectors. More events and structure is present in the Allen dataset. However, the stimuli were different (see Allen whitepapers). (**D**) Example events from Allen dataset. (**E**) Core neurons (green) were evenly distributed along events. (**F**) Distributions of pattern reproducibility for the MICrONS phase 1 dataset (black) and each (*N*=32) Allen Brain Observatory (gray). The MICrONS distribution does not qualitatively differ from those from the Allen Brain Observatory. Note that Allen recordings are on average three times longer than the MICrONS, resulting in a reduced count for the latter.

**Fig. S3.**
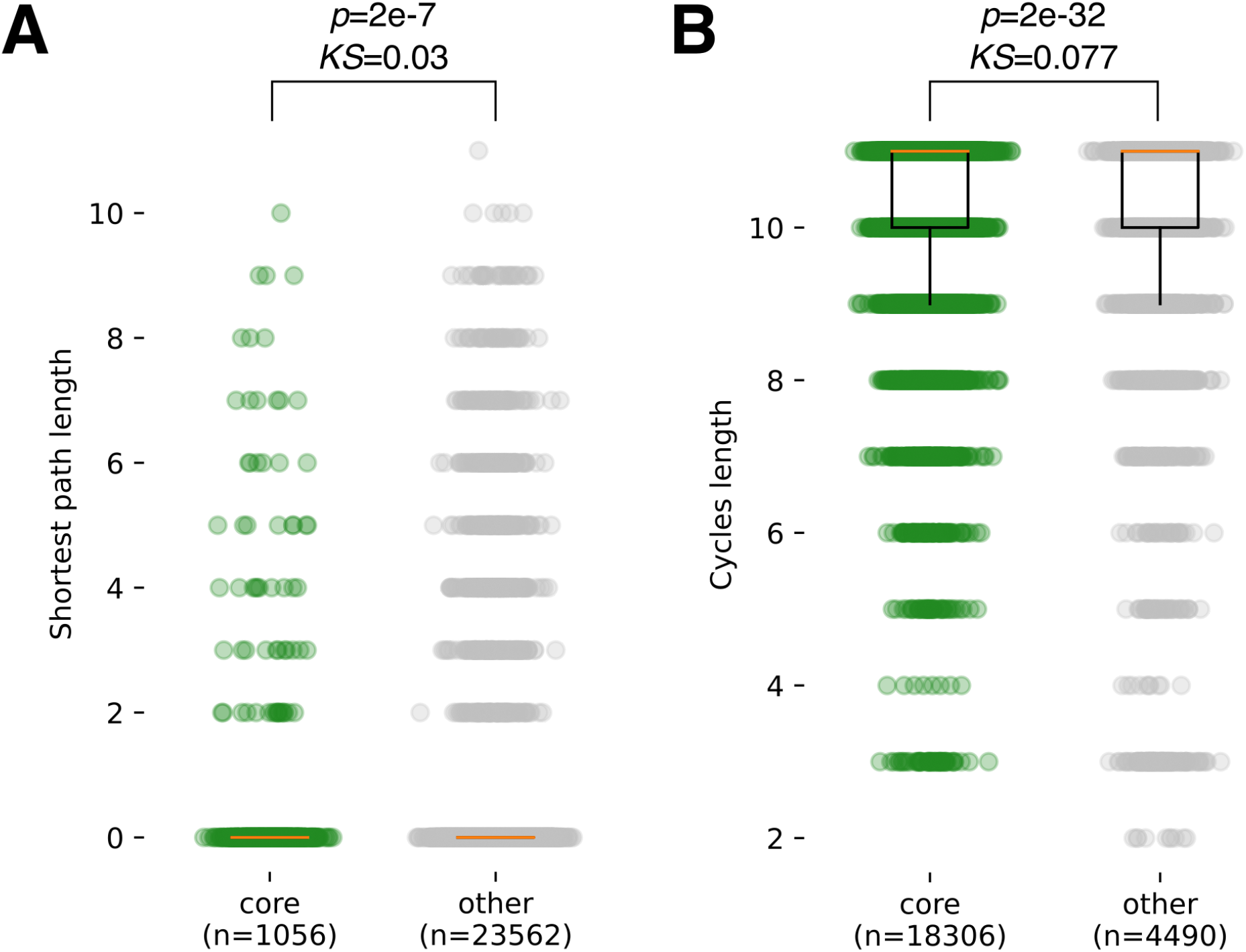
Cores are not recursively connected via multi-synaptic feedback. Even if cores were not directly connected, they could be highly connected via secondary paths, abundant enough to ensure that they are pattern completion units in traditional attractor network terms. (**A**) Core-to-core shortest paths were neither more nor shorter than other-to-other paths (0.27±1.19 vs 0.16±0.92 length, Kruskal-Wallis t=26.4 p=2.7e-07, *KS=0.03*). (**B**) Core neurons could be part of looped paths, circling back to them, and providing a weak form of recursion. Core-based cycles were more (green, 18306) compared to other-based cycles (gray, 4490). But core and non-core cycles had on average the same length (10.34±1.07 and 9.93±1.76 connections, with minimal effect size KS=0.077), same as the network diameter (*d*=10).

**Fig. S4.**
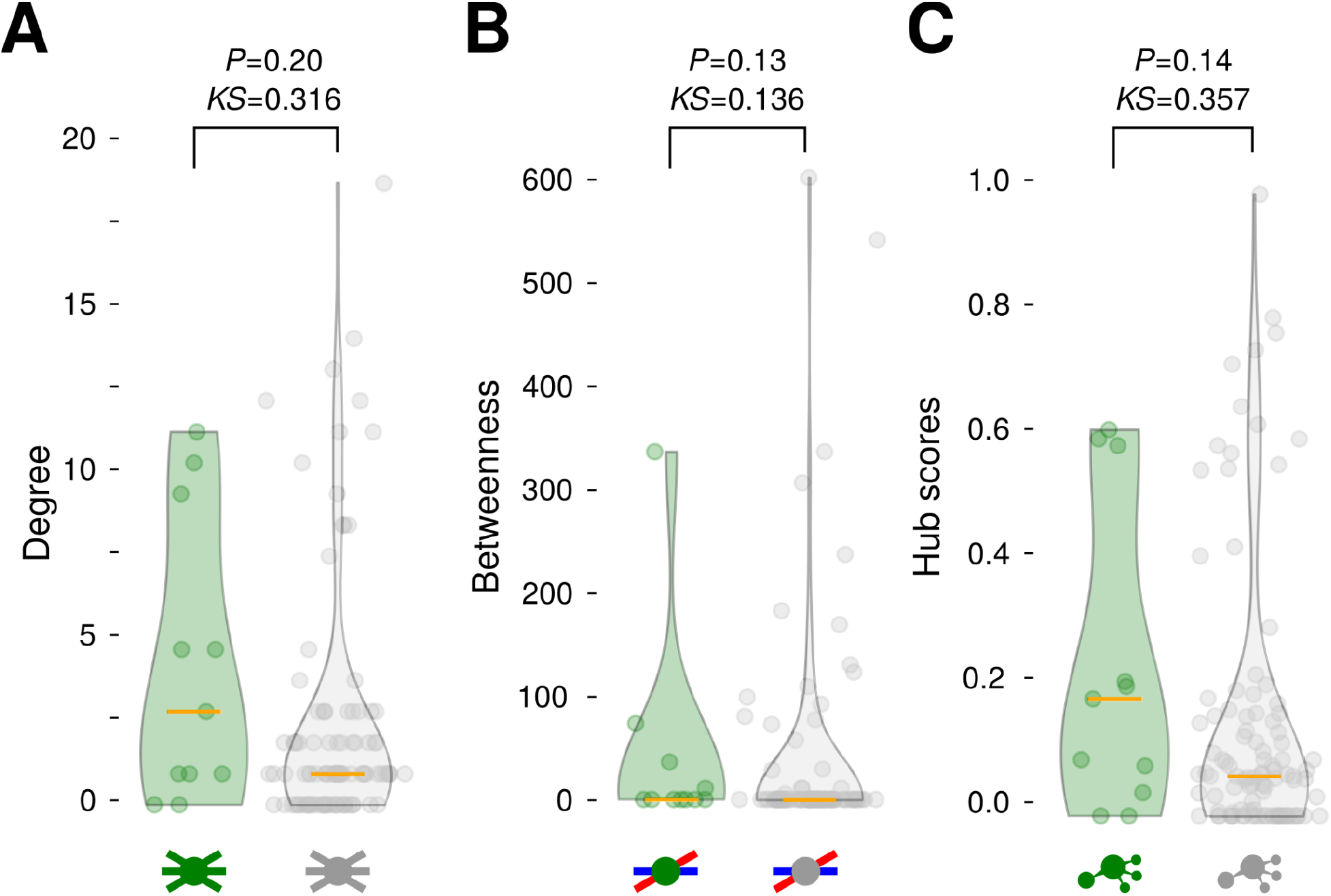
Simple centrality measures show no difference between cores and other neurons. (**A**) The degree – number of connections per neuron – of cores was (not significantly) higher than other neurons. (**B**) The betweenness – number of shortest paths between any two nodes passing by a considered node – of cores was not significantly different from other neurons. (**C**) The hub score – the weighted number of outward connections of a node that point to central nodes – of cores was (not significantly) higher than other neurons.

**Fig. S5.**
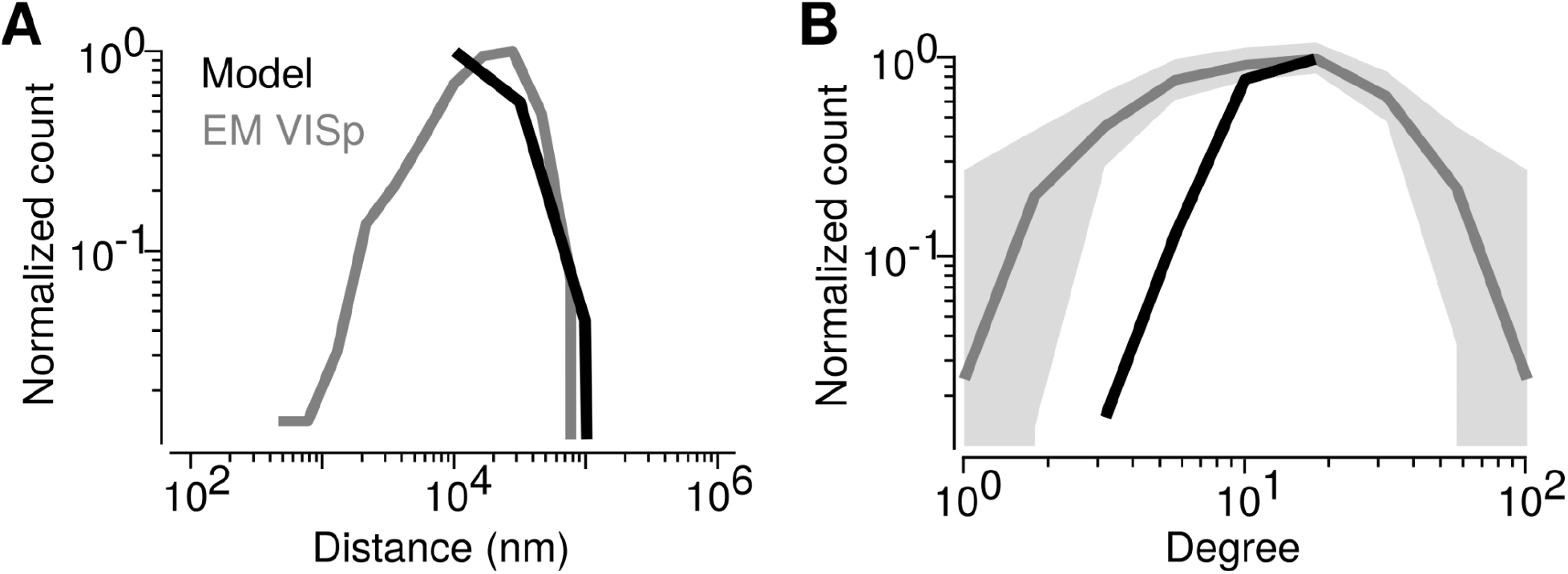
Comparison of the MICrONS and model basic network properties. (**A**). (**B**) Normalized degree of degrees MICrONS degree max is 75 (gray), which we imposed as maximal value in all models (black). For small distance-dependent connectivity ranges (from 22 to 42 μm) the number of actual connections was lower than 75 because of the cell density. In these models, the cells were connected with all neighbours.

**Fig. S6.**
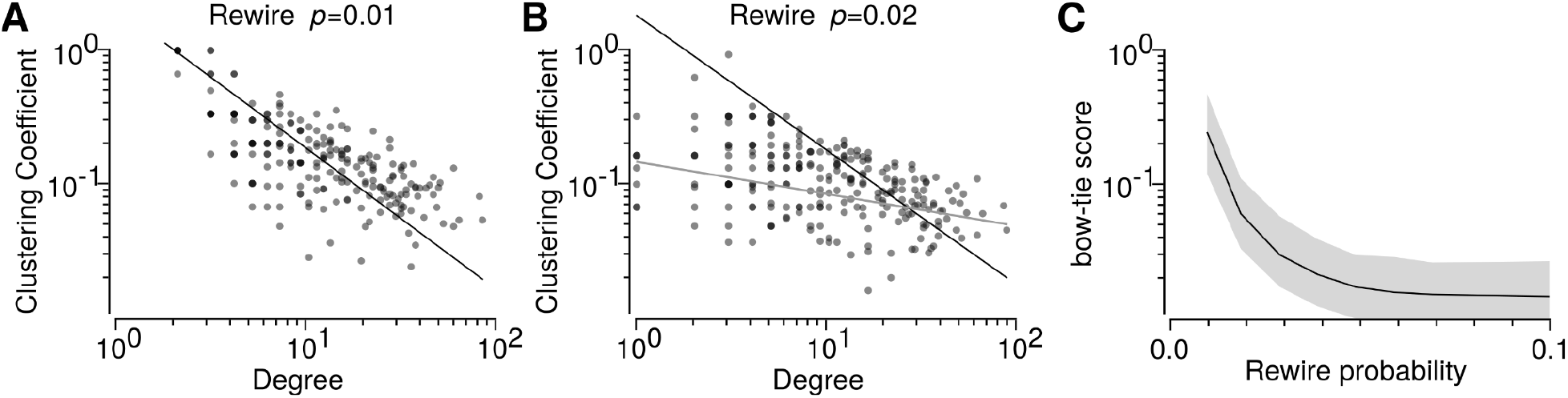
Bow-tie evidence in the MICrONS dataset and overlap between dynamical and structural cores. **A** and **B**. Example hierarchical modularity resulting from rewiring the MICrONS graph. In **A**, rewiring the graph edges with probability *p*=0.01 results in a hierarchically modular relationship as that observed in the data. In **B**, already at *p*=0.02 the relationship deteriorates (fitting curve is kept identical to the data in both panels, with a). **C**. The bow-tie score (ratio of modules with clearly identifiable submodules converging-diverging from a central module over the total number of modules) rapidly deteriorates for increasing rewiring probabilities.

**Fig. S7.**
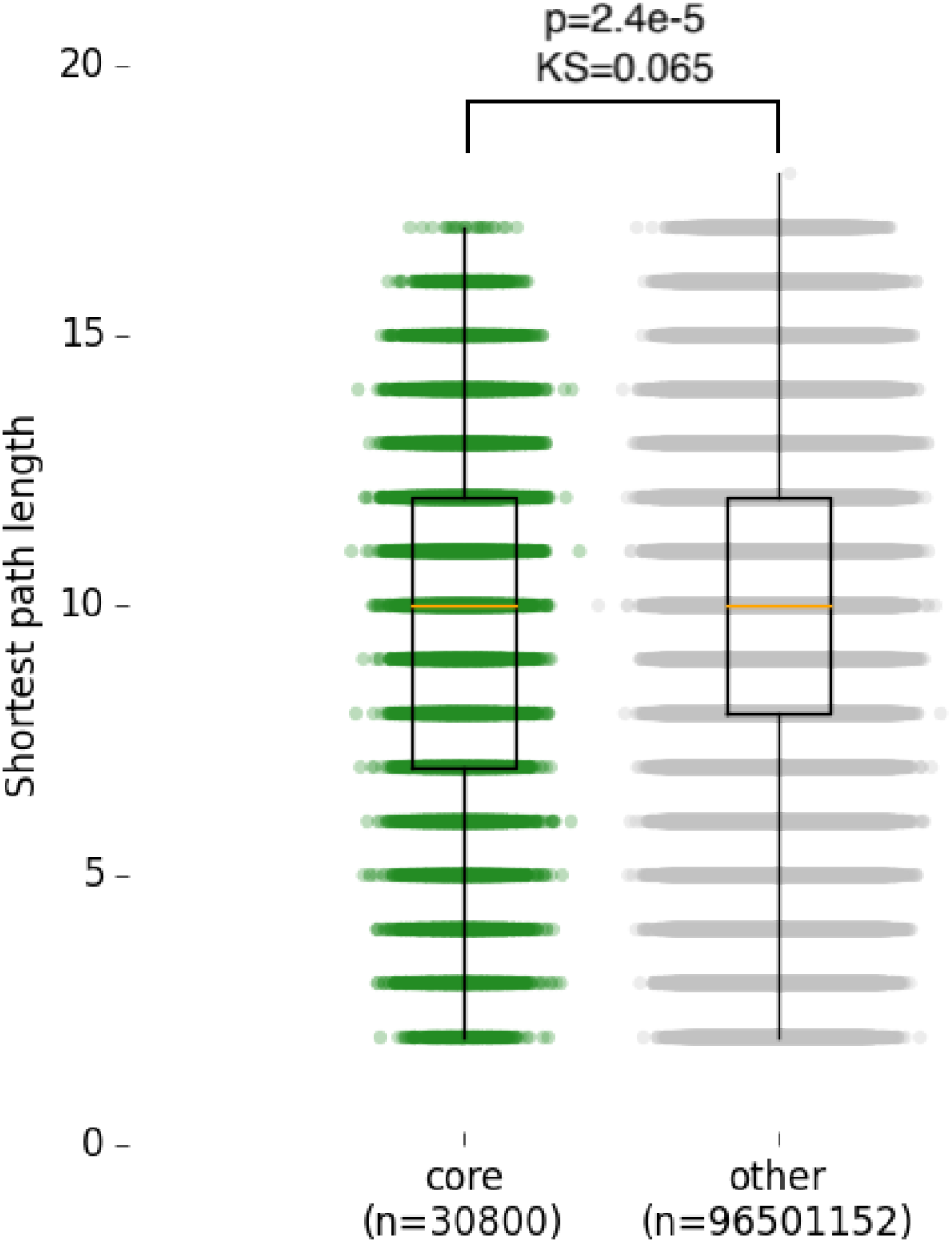
The shortest paths between model cores and others were not different. Even if cores were not directly connected, they could be highly connected via secondary paths, abundant enough to ensure that they are pattern completion units in traditional attractor network terms. To evaluate this aspect also in the model as in the data (see Fig. S2), we measured the number and length of shortest paths between cores and compared them with those made by others. We found that the shortest paths between cores were significantly shorter compared to those between others (cores: 9.75±3.36 vs others: 10.04±2.98 length), overall with negligible effect size (*KS* = 0.065). A direct comparison of cycles in the model is computationally infeasible.

**Fig. S8.**
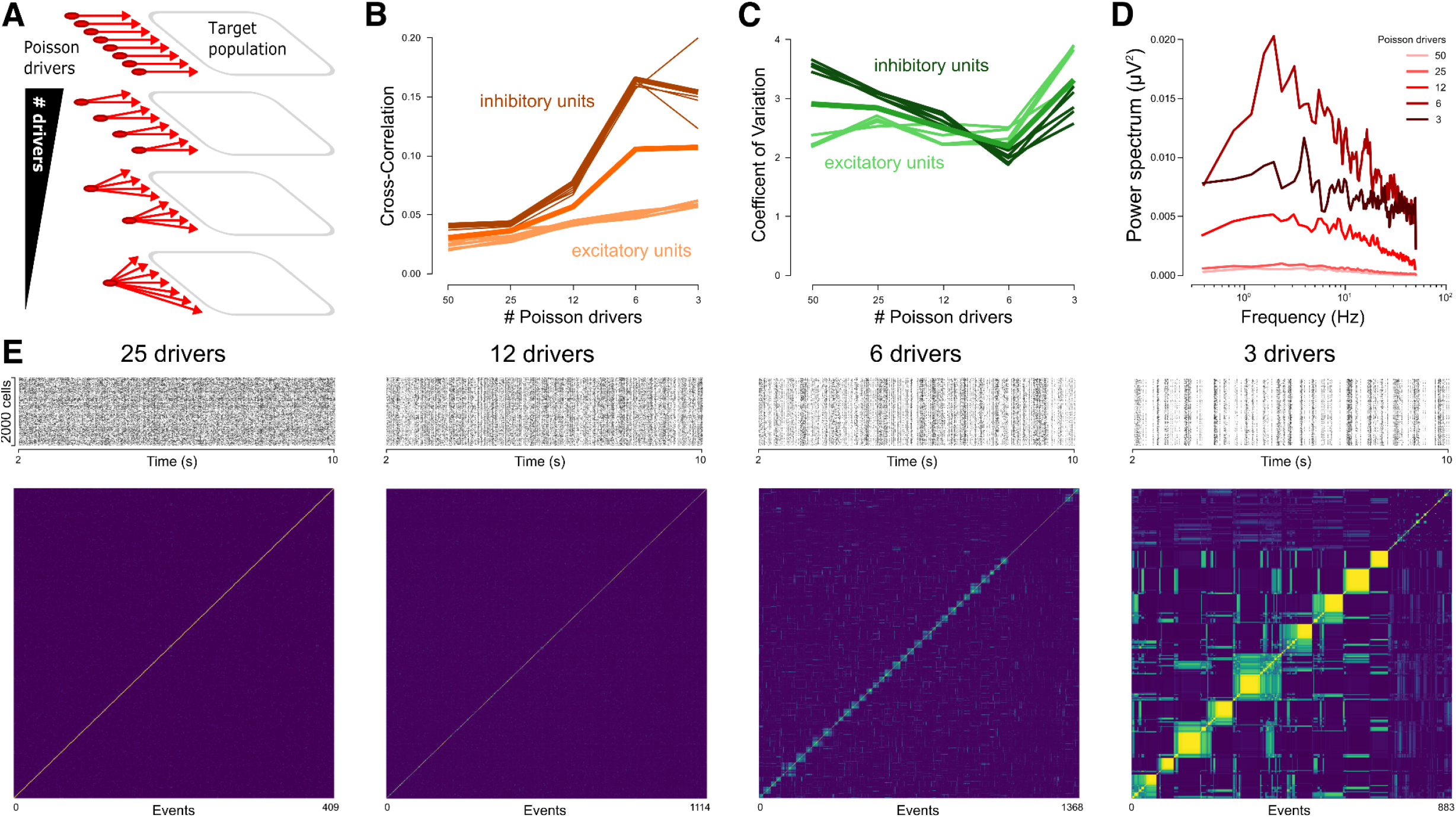
Increasing input correlations rescues population event reproducibility. (**A**) For the same target population (right), we systematically reduced the number of Poisson input drivers (red disks and arrows), to explore how the increase in input correlations affected the correlations between population events. (**B**) Reducing the number of Poisson input drivers (n=[50, 25, 12, 6, 3]) increased the population cross-correlation of spikes toward values characteristic of oscillating regimes (from CC=0.035 at n=50, to CC=0.11 at n=3). (**D**) Reducing the number of Poisson input drivers increased the power for slow oscillations in the Fourier spectrums of the firing rates. (**E**) Four 10 sec spike rasters for 2000 example cells, with their correlation matrices, show the changes in firing regimes responsible for the increase in population events correlation. The first correlation matrix on the left shows the absence of reproducibility. In the right correlation matrix, the off-diagonal correlations show the presence of correlations between population events, resulting in higher values of reproducibility.

**Fig. S9.**
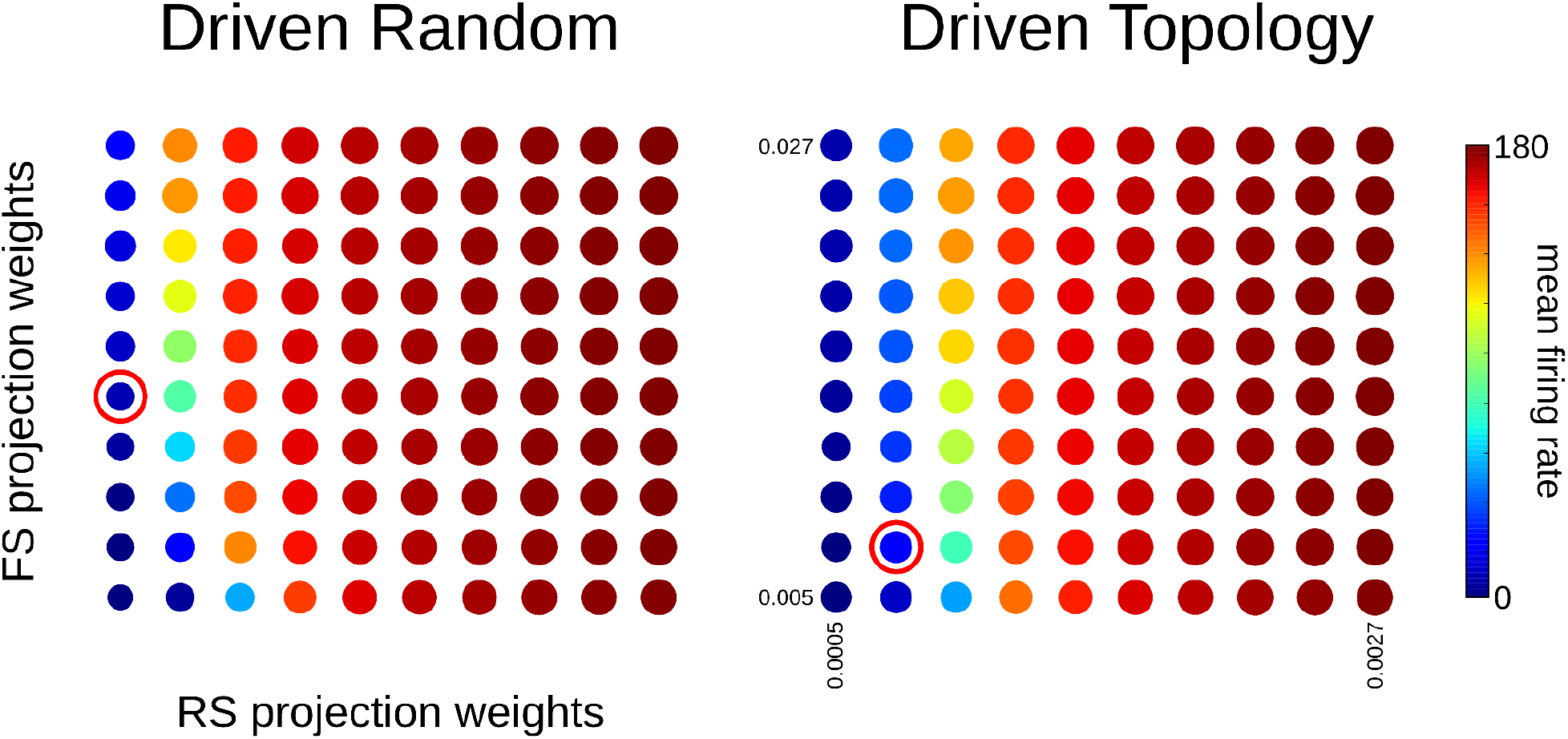
The model is robust with respect to changes in synaptic parameter space. We explored the parameter space made by the relationship between regular spiking (RS) and fast spiking (FS) synaptic conductance change triggered by incoming spikes, for both the randomly connected (*left*) and topologically connected (*right*), for what concerns the stability of firing regime. Many couples of synaptic parameters produced stable firing rates too high to be comparable with the 2-photon activities (from 80 spikes/s on, from cyan to red, see colorbar on the right). We therefore further tested several couples below 80 spikes/s and settled on 5-10 spikes/s (marked by the red circle).

